# Comparative genomics of four *Coptis* species provides insights into rhizome development and alkaloid biosynthesis

**DOI:** 10.64898/2026.11.10.765818

**Authors:** Mingjiang Huang, Yingchun Zhang, Lei Pang, Mengguo Ran, Shengyong Chen, Yufeng Ruan, Wanqing Liu, Tingrui Peng, Tao Yu, Jian Qin, Difeng Tan, Xuefeng Yu, Xin Tang, Guanglou Nie, Pingan Peng, Xing Zhao, Xuegang Li, Hang Ma, Xiaoli Ye

## Abstract

*Coptis* species belong to the Ranunculaceae family and have important pharmacological activity because their rhizomes are rich in isoquinoline alkaloids. However, the genome evolution and species relationships of *Coptis* remain poorly understood. Here, we report high-quality chromosome-level genome assemblies of four representative *Coptis* species (*C. chinensis*, *C. deltoidea*, *C. omeiensis*, and *C. teeta*). We found that the evolution of this group was strongly affected by geological events, especially the uplift of the Tibetan Plateau and the collision between the Indian and Eurasian plates. The triploid *C. deltoidea* exhibited striking B subgenome expression dominance, while *C. omeiensis* showed extensive tandem duplication of *NLR* disease-resistance genes. Rhizome branching and swelling were closely associated with the activation of brassinosteroid biosynthesis and signaling pathways. Key gene families involved in alkaloid biosynthesis were significantly expanded in *Coptis* and were mainly highly expressed in the rhizome. Multiple transcription factor families, including MYB, bHLH, and ERF, were identified as candidate regulatory factors associated with alkaloid biosynthesis. We further constructed the first public multi-omics database for *Coptis*. This study provides key genomic resources for understanding *Coptis* evolution, adaptation, and molecular breeding.

## Introduction

*Coptis* (Ranunculaceae), a genus of perennial medicinal plants, has significant antibacterial, anti-inflammatory, and gastrointestinal functional regulatory effects because of the abundance of isoquinoline alkaloids such as berberine and coptisine in its roots and rhizomes^1-3^. It is known as the “golden herb” in traditional Chinese medicine. Currently, three species of *Coptis* are included in the Chinese Pharmacopoeia, namely, *C. chinensis* from Chongqing and Hubei, *C. deltoidea* from Sichuan, and *C. teeta* from Yunnan and Xizang, which are commonly known as ‘Weilian’, ‘Yalian’ and ‘Yunlian’, respectively (Fig. 1A). In addition, some species that are not included in the pharmacopoeia, such as *C. omeiensis*, *C. japonica*, and *C. chinensis* var. *brevisepala*, also have important medicinal value^4-7^.

**Fig 1.**
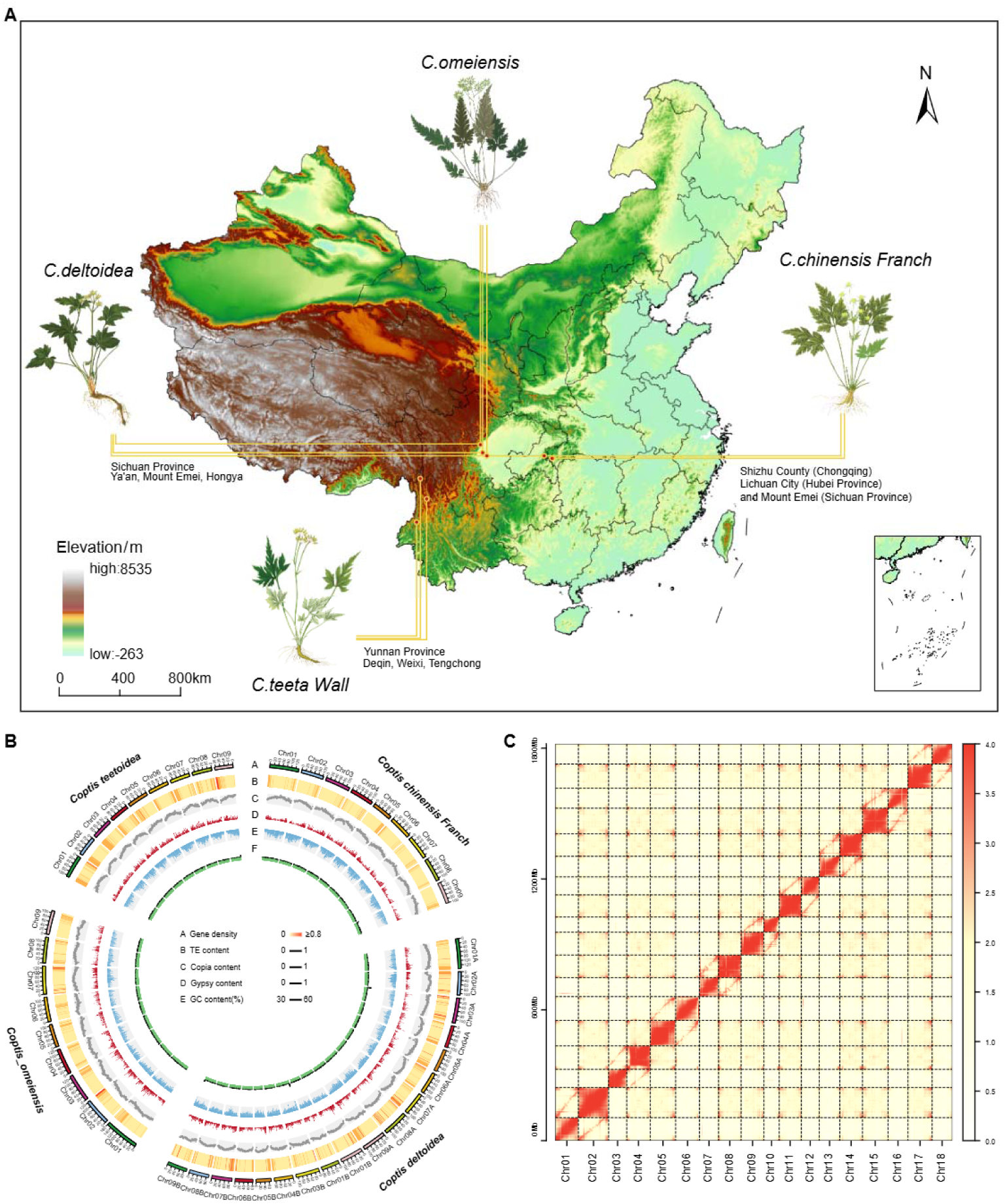
Chromosome-level genome assemblies and features of four *Coptis* species. (A) Native ranges and sampling localities of four *Coptis* species in China overlaid on a digital elevation map.(B) Circos view of chromosome ideograms with genomic feature tracks (e.g., gene density, GC/repeat content) and intergenomic synteny links among species/subgenomes.(C) Hi-C contact heatmap of *C. deltoidea* showing chromosome-level scaffolding (18 pseudochromosomes).

Different species of *Coptis* exhibit significant differences in growth habits, traits, and alkaloid contents^4,5,7^. For example, *C. chinensis* is suited for growth at altitudes of 1200–1800 metres, *C. deltoidea* grows at altitudes of 1800–2500 metres, and ***C. teeta*** grows at altitudes above 1900 metres. In terms of plant morphology, *C. chinensis* has multiple branching rhizomes, forming clusters resembling chicken feet, commonly known as ‘Jizhualian’ (Fig. 1A). The rhizomes of *C. deltoidea* and *C. teeta* are single branched and relatively slender. Therefore, the biomass of the rhizomes of *C. chinensis* is greater than that of *C. deltoidea* and *C. teeta*, making it the main source of *Coptis* on the market. Karyotype analysis has also revealed differences in ploidy among different *Coptis* species; while most species are diploid (2n = 2x = 18), triploids, such as *C. deltoidea* (2n = 3x = 27)^8^, have also been detected. These marked differences in ploidy, morphology, rhizome traits, and altitudinal distribution among closely related *Coptis* species provide an ideal natural system for investigating how geological events, subgenome evolution, and hormonal regulation contribute to species diversification, environmental adaptation, and medicinal trait formation.

To date, several chromosome-level *Coptis* genomes have been assembled. For example, Liu et al^9^ assembled a high-precision chromosome-level genome of *C. chinensis* (∼936.6 Mb) and revealed that the functional diversity of a key CYP719 gene involved in the berberine biosynthesis pathway may play a crucial role in the diversification of other berberine-related alkaloids. Wang et al^10^ successfully assembled and annotated the haplotype reference genome of *C. teeta* and reported numerous structural alterations within nine homologous chromosome pairs, which may lead to aberrant synapsis and irregular chromosomal segregation, thereby contributing to the sustained preservation of clonal propagation. However, it is difficult to analyse key issues such as the evolutionary history and rhizome branching mechanisms of *Coptis* species through studies of a single species.

Assembling multiple high-quality genomes of *Coptis* species will benefit research on species evolution and genetic breeding in this genus. Therefore, in this study, we assembled high-quality chromosome-level genomes of four representative *Coptis* species with important medicinal value, namely, *C. chinensis*, *C. deltoidea*, *C. omeiensis*, and *C. teeta*. Through integrated multiomics analysis, we systematically revealed the evolutionary history, environmental adaptation mechanisms and elongation and expansion mechanisms of the rhizome. Furthermore, we established a comprehensive multiomics database for *Coptis*. Our study provides key theoretical support for the conservation and development of *Coptis* germplasm resources.

## Results

### Genome assembly and annotation of C. chinensis, C. deltoidea, C. omeiensis and C. teeta

Based on k-mer analysis of DNBSEQ short reads with an average coverage of 78.5–201.9×, the estimated genome size of the four *Coptis* species was 973.1–1782.8 Mb (Supplementary Fig.1 and Supplementary Table 1). Using PacBio long reads with an average coverage of 43×, the four genomes were de novo assembled into 155 to 761 contigs and polished, yielding minimum and maximum contig N50 values of 4.59 Mb and 13.5 Mb, respectively (Supplementary Table 2-3). The assembly result is highly consistent with the genome size estimated by *k-mer* analysis, verifying the accuracy of the assembly. Based on Hi-C sequencing data, the total assembly sizes of the four *Coptis* genomes ranged from 914.85 Mb to 1,922.15 Mb, with the largest corresponding to the only triploid species, *C. deltoidea*. The scaffold N50 values ranged from 92.03 Mb to 181.89 Mb, with an average of 125.30 Mb (Fig. 1B and 1C, Supplementary Fig.2 and Supplementary Table 3).

To evaluate the assembly quality of the four genomes, we compared the DNBSEQ short reads with the assembled genomes and obtained an average mapping rate of 95.00%. BUSCO evaluation revealed that the assembly integrity exceeded 95.00%. These results indicated the high continuity and integrity of the four genomes (Supplementary Fig.3). A combination of homologous and de novo evidence indicated that the proportion of repetitive sequences in the four genomes was 63.89–72.53%, with the highest proportion of long terminal repeat sequences (LTRs) (46.37–53.64%). These LTRs belong mainly to the Gypsy (38.31–45.49%) and Copia (6.65–7.25%) families, which is consistent with the transposon burst characteristics commonly found in Ranunculaceae plants (Supplementary Fig.4 and Supplementary Table 4)^11-13^. Through a combination of de novo, homology, and transcript-based prediction results, 51224 protein-coding genes were predicted in the four assembled genomes. BUSCO analysis indicated that the annotation integrity of the four genomes exceeded 95.00% (Supplementary Fig.5). The predicted protein-coding genes were subsequently compared with protein sequences in six databases, namely, Nr, SwissProt, KEGG, COG, TrEMBL and InterPro, and an average of 85% of genes in each genome were assigned putative functional annotations. Intriguingly, although the four *Coptis* species shared similar genomic architectures, *C. teeta exhibited distinct structural divergence. Statistical analysis revealed that both exon and intron lengths in C. teeta* were significantly greater than those in *C. chinensis*, *C. deltoidea*, and *C. omeiensis* (Supplementary Fig. 6).

### Analysis of gene families and phylogenetic relationships

To elucidate the phylogenetic relationships among the four *Coptis* species and other angiosperms, we performed a cluster analysis of genes from these four *Coptis* species and five other plant species, including *Phellodendron amurense*, *Papaver somniferum*, *Aquilegia vulgaris*, *Kingdonia uniflora* and *Oryza sativa*. In total, 406,865 genes were clustered into 35,193 gene families, of which 77 single-copy gene families were shared by all nine species. Notably, 26 homologous groups contained 10,446 genes unique to the *Coptis* species (Fig.2B,2C, Supplementary Fig.7-10 and Supplementary Table 6-7). Based on the time-scaled phylogenetic tree, we found that *Coptis* and *A. vulgaris* diverged approximately 73.9 Mya (million years ago), which is consistent with previous reports^9^. Additionally, our analysis revealed that the first divergence within the *Coptis* lineage occurred approximately 18 Mya, with the A subgenome of *C. deltoidea* differentiating from the *C. omeiensis* genome and B subgenome of *C. deltoidea* occurring 13.6 Mya. The *C. omeiensis* and B subgenomes of *C. deltoidea* subsequently diverged 7.3 Mya (Fig. 2A). Interestingly, the divergence time between *Coptis* and *A. vulgaris* falls within a broad interval that overlaps with major India–Eurasia collision-related geological reorganization. The two internal divergence events within the *Coptis* lineage broadly coincide with key geological and climatic events in the early to middle Miocene (23–15 Mya and 10–7 Mya) in the Qinghai–Tibet Plateau–Himalayan–Hengduan Mountains region, including the initial rapid uplift of the Tibetan Plateau and the concurrent intensification of the Asian monsoon^14,15^.

**Fig 2.**
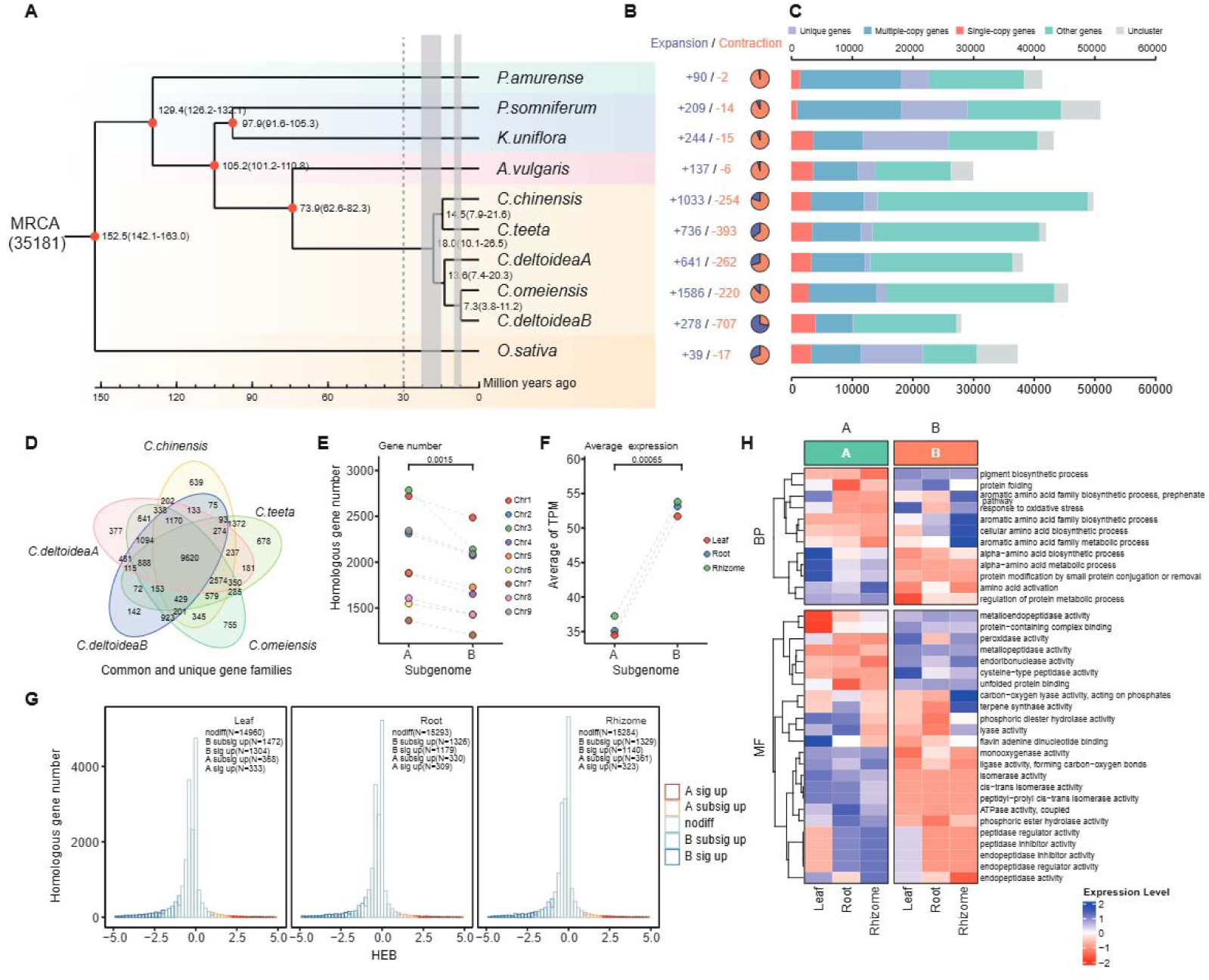
Genome evolution and subgenome divergence in *Coptis*. (A) Chronogram of *Coptis* and related taxa with divergence times (Mya); red dots indicate calibrated nodes, and bars denote 95% HPD intervals.(B) Numbers of gene family expansions (+) and contractions (–) along each lineage.(C) Gene repertoire composition per genome, showing counts of unique, multicopy, single-copy and other genes. The species included *C. chinensis*, *C. teeta*, *C. omeiensis*, and *C. deltoidea* (A/B subgenomes), with representative outgroups.(D) Venn diagram of shared and unique gene families across five genomes/subgenomes (*C. chinensis*, *C. teeta*, *C. omeiensis*, *C. deltoidea*-A, and *C. deltoidea*-B).(E) Comparison of homoeolog numbers between subgenomes A and B (each point = one homoeologous chromosome pair).(F) Comparison of average gene expression (TPM) between subgenomes A and B.(G) Distributions of homoeolog expression bias (HEB) in leaves, roots and rhizomes; red/blue indicate A-/B-biased genes, grey indicates not significant genes.(H) GO enrichment heatmaps (BP, MF) for A- and B-biased genes across tissues; colour denotes the relative expression/enrichment level (legend).

We further conducted positive selection analysis on the background branch of *C. chinensis* and identified a total of 13 positively selected genes (Supplementary Table 8). Subsequently, functional enrichment analyses were performed for the expanded/contracted gene families and the positively selected gene families. The expanded gene families were predominantly enriched in biological processes associated with homeostatic regulation, particularly ion and metal ion homeostasis, which are critical for maintaining cellular stability and supporting continuous cell division and proliferation (Supplementary Fig.11 and Supplementary Table 9-10). Notably, we identified the *BR6ox1* gene, a key enzyme involved in brassinosteroid (BR) biosynthesis^16,17^, within the expanded families. Given the established role of BRs in promoting cell division and elongation, the expansion of *BR6ox1-like genes* is likely to have contributed to enhanced BR production and signalling, thereby facilitating rhizome branching and development in *C. chinensis*. In contrast, the positively selected genes were significantly enriched in RNA processing, DNA mismatch repair, protein lipidation, and GPI anchor metabolic processes, functions that are closely related to transcriptional regulation, genomic stability, and signal transduction (Supplementary Table 11-12).

### Genomic and ecological analysis of the expression advantage of the B subgenome in *C. deltoidea*

Due to the AAB genotype of *C. deltoidea*, the number of homologous genes in the A subgenome is significantly greater than that in the B subgenome (Fig. 2E). However, an analysis of subgenome expression patterns revealed that the total gene expression in the B subgenome was notably higher than in the A subgenome (Fig. 2F and Supplementary Fig.13B). Further investigation revealed that the average gene expression and the gene expression across different tissues in the B subgenome were significantly elevated compared with those in the A subgenome, supporting a marked expression advantage of the B subgenome (Fig. 2F and 2G, Supplementary Fig.13C). We subsequently conducted an enrichment analysis of the dominant genes in the A/B subgenomes and reported that the A subgenome exhibited significant expression advantages in processes such as pigment synthesis and protein and amino acid metabolism (Supplementary Fig.14). These expression patterns may contribute to growth, development, environmental adaptation, and metabolic differentiation in C. deltoidea(Fig.2H). Conversely, the B subgenome presented higher expression levels of genes related to enzyme activity, the oxidative stress response, and protein binding and transport (Fig.2H). These findings suggest that the expression advantage of the B subgenome plays a crucial role in enhancing the metabolic capacity, antioxidant stress resilience, and maintenance of protein homeostasis of plants.

### Karyotype reconstruction and adaptive evolution of *Coptis*

The emergence of the AAB genomic constitution coincides with major geological and climatic events during the early to middle Miocene, particularly in the Qinghai–Tibet Plateau–Himalayan–Hengduan Mountains region (23–15 Mya and 10–7 Mya)^14,15^. This period marked a critical point in plant evolution driven by climatic and ecological shifts. The geological transformations during this time may have played an important role in shaping the genomic architecture of *C. deltoidea*. Specifically, environmental changes may promote adaptive variation and its retention within populations. Among these, structural variations in chromosomes may be associated with such pressures and contribute to the formation of lineage-specific traits. To further investigate this hypothesis, we performed ancestral karyotype reconstruction. The results revealed a clear contrast between subgenomes: while the B subgenome remained largely conserved with no detectable chromosomal rearrangements, the A subgenome experienced two major chromosomal breakage events (Fig. 3 and Supplementary Fig. 15). These breakpoints were identified on specific chromosomes and are supported by the presence of truncated gene structures in multiple species (e.g., *Ccs01G000600*, *Cte01G052700*, and *Com09G059600*), with break positions spanning distinct genomic intervals, indicating conserved yet lineage-specific structural disruptions.

**Fig 3.**
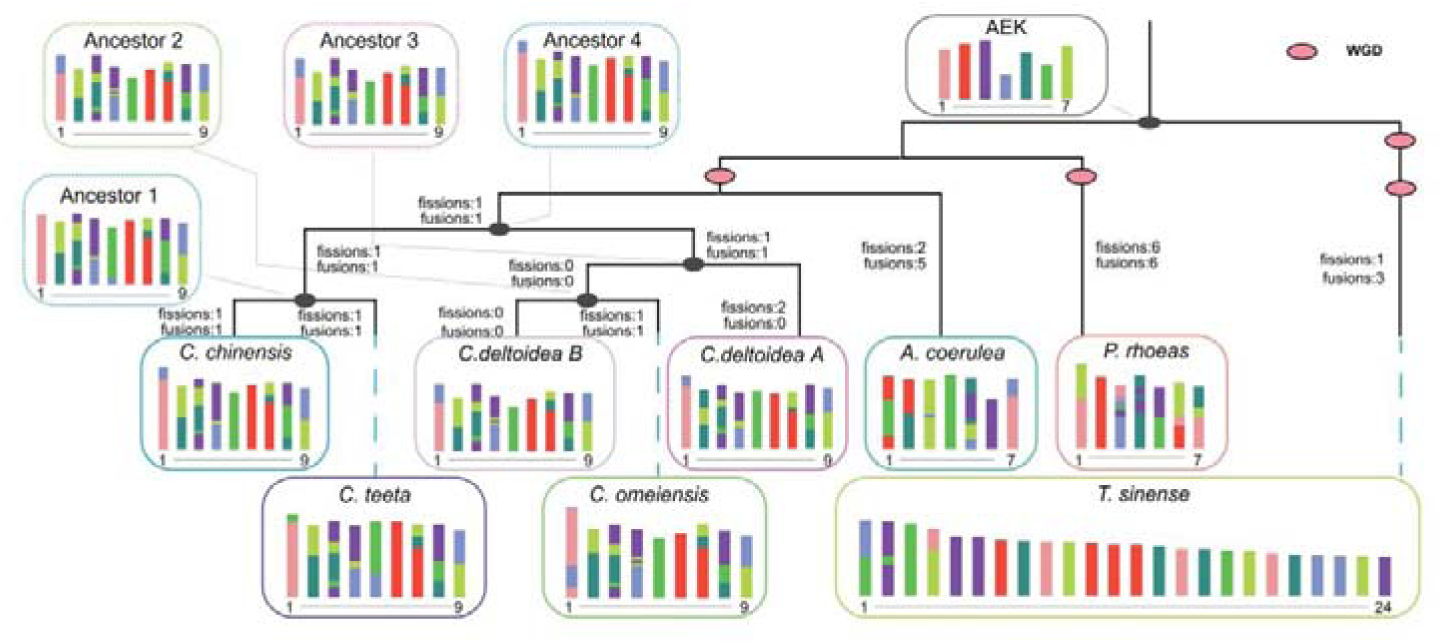
Karyotype evolution of *Coptis*. (A) Reconstruction of the rearrangement history from the ancestral eudicot karyotype (AEK) to extant *Coptis* genomes, indicating inferred whole-genome duplications (WGD) and lineage-specific fissions, fusions and translocations that produced the A and B subgenomes of *C. deltoidea* as well as the diploids; *Aquilegia coerulea*, *Papaver rhoeas* and *T. sinense* were used as outgroups

In contrast, the structural stability of the B subgenome throughout these events may have conferred adaptive advantages by preserving essential gene functions and regulatory networks. This genomic conservation likely underlies its relative expression dominance in certain metabolic pathways critical for plant survival.

### Identification and expression analysis of *NLR* genes

In response to extreme environmental pressures, plants typically enhance their survival fitness by boosting disease resistance, a process that is frequently accompanied by the expansion and functional diversification of resistance-related gene families^18-20^. In this study, we systematically identified nucleotide-binding site leucine-rich repeat (*NLR*) genes across *Coptis* species at the genome-wide level (Fig. 4A and Supplementary Fig.16). The results showed that the number of *NLR* genes in the A subgenomes of *C. deltoidea* and *C. omeiensis* was significantly higher than that in other genomes. Meanwhile, no TIR-NLR (*TNL*) genes were detected in any of the examined *Coptis* species, suggesting that this lineage may have undergone lineage-specific loss of *TNL* genes during the long-term evolutionary process.

**Fig. 4.**
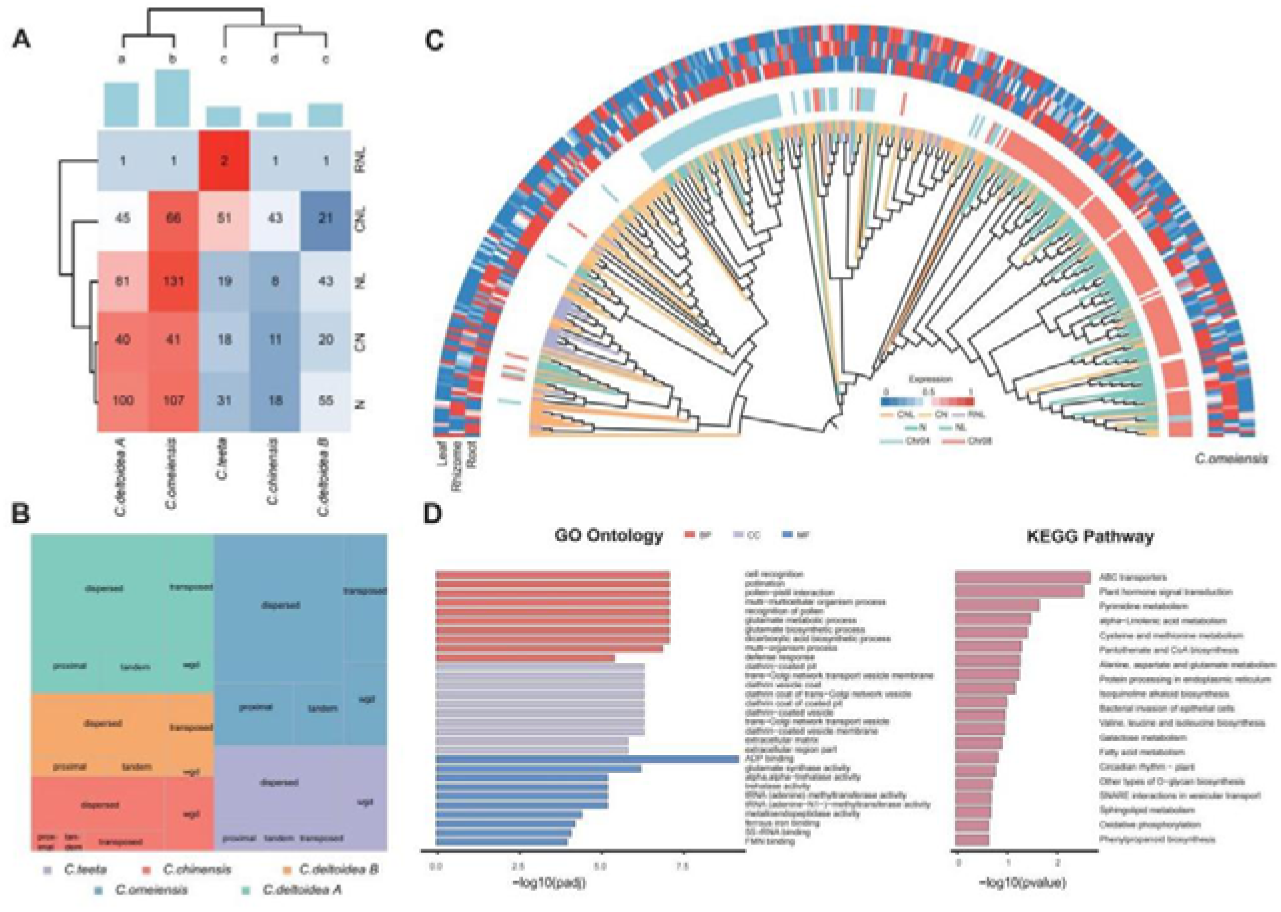
*NLR* gene diversification in *Coptis*. (A) Heatmap with hierarchical clustering of NLR domain category sizes across *C. chinensis*, *C. teeta*, *C. omeiensis*, *C. deltoidea*-A and *C. deltoidea*-B.(B) Genome-wide distribution of duplication modes for NLR genes in the five species/subgenomes (WGD, tandem, proximal, dispersed and transposed).(C) Maximum-likelihood phylogeny of NLR proteins (*C. omeiensis*), with outer tracks indicating species provenance and duplication mode.(**D**) Gene Ontology (GO) terms and KEGG pathways enriched among *NLR*-related genes/domains in *C. omeiensis*.

Further analysis of *NLR* gene duplication events in the *Coptis* genomes revealed that *C. omeiensis* had the highest number of gene duplication events, followed by the A subgenome of *C. deltoidea* (Fig. 4B and Supplementary Fig.17). These results indicate that these two genomes/subgenomes may have contributed to adaptation to the extreme environmental pressures generated during geological changes through the amplification and duplication of key resistance genes. To further investigate the evolutionary dynamics of the *NLR* gene family, we constructed a phylogenetic tree for *C. omeiensis*, which shows the most pronounced expansion of *NLR* genes. The results indicate that different types of *NLR* genes are distributed across multiple clades with clear clustering patterns (Fig. 4C). Integrated analysis with transcriptomic expression data revealed that these genes were predominantly highly expressed in root tissues. This tissue-preferential expression pattern suggests that as perennial herbaceous plants, the rhizomes of *Coptis* are continuously exposed to the soil environment, and these *NLR* genes may play a critical role in defending against soil-borne pathogen infection. Furthermore, functional enrichment analysis of the expanded genes in *C. omeiensis* showed that these genes were significantly enriched in biological processes and signaling pathways related to disease resistance and reproduction, including multiple Gene Ontology (GO) terms and Kyoto Encyclopedia of Genes and Genomes (KEGG) pathways (Fig. 4D), further supporting their important functions in environmental adaptation.

Collectively, our findings suggest that *Coptis* species have evolved adaptive mechanisms to cope with complex environmental pressures through the expansion, duplication, and expression divergence of the *NLR* gene family, a process that may have been associated with historical geological and ecological changes.

### Molecular mechanism analysis of high-yield traits in *C. chinensis* rhizomes

Comparative genomic analyses revealed that genes under positive selection in *C. chinensi*s are significantly enriched in functional categories associated with cell elongation and expansion. Notably, gene family expansions specific to *C. chinensi*s were predominantly involved in BR biosynthesis and signalling pathways. Given the well-established roles of BRs in promoting cell division and expansion during root and rhizome development^17,21,22^, these findings suggest that BRs may play crucial regulatory roles in the growth and development of *C. chinensi*s. Phenotypic observations further support this molecular evidence (Fig. 5A). Compared with the other three *Coptis* species (*C. deltoidea*, *C. omeiensis*, and *C. teeta*), *C. chinensis* presents a markedly greater number of rhizome branches and significantly elongated and swollen branch nodes, ultimately contributing to a substantial increase in rhizome yield (Fig.5B).

**Fig. 5.**
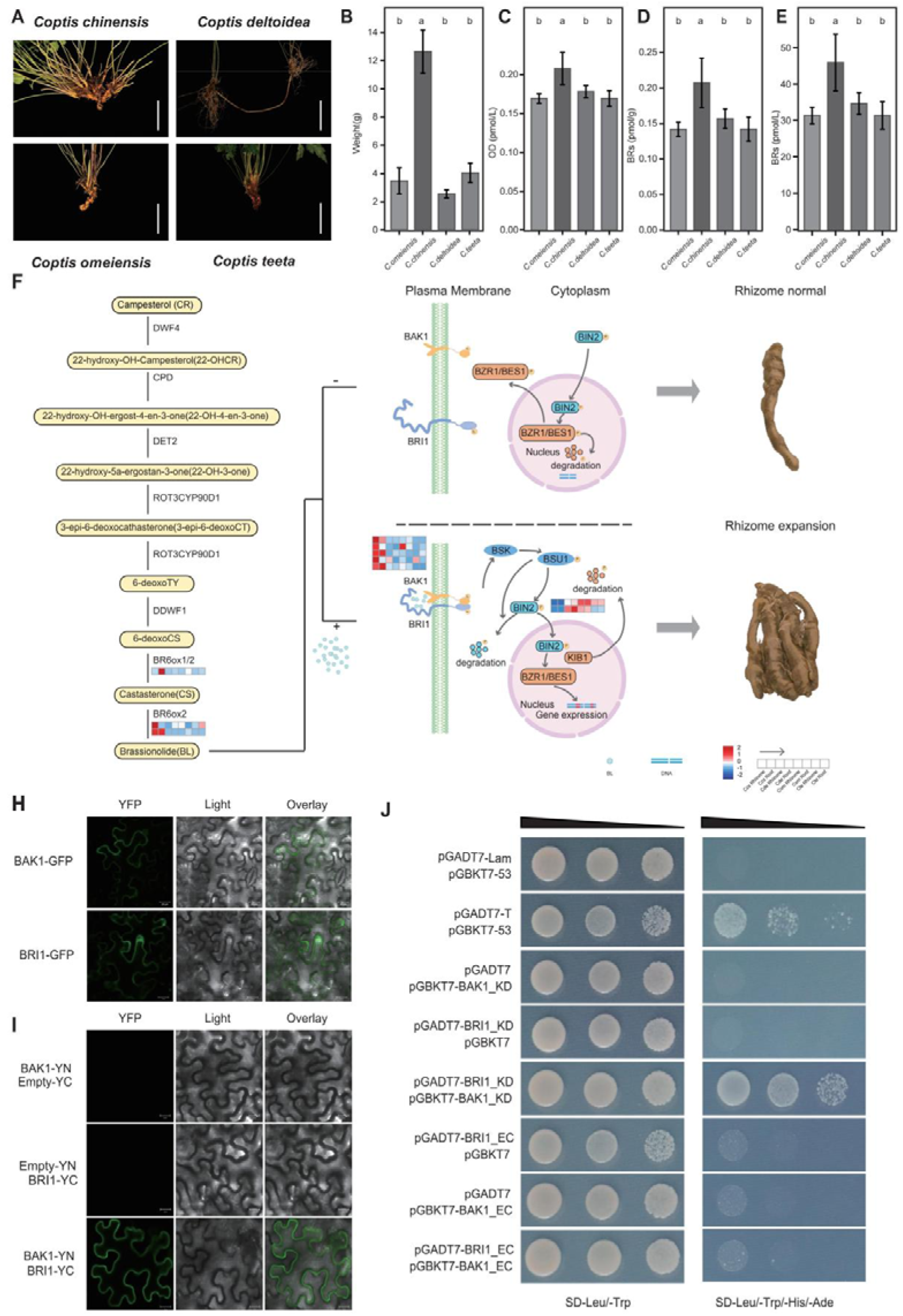
BR pathway activation is associated with rhizome branching and swelling in *C*. *chinensis*. (A) Representative phenotypes of four species (*C. chinensis*, *C. deltoidea*, *C. omeiensis*, and *C. teeta*).(B) Rhizome weight of the four species.(C–E) Endogenous BR-related metabolite levels by species; letters above bars indicate Tukey groupings.(F) BR biosynthetic pathway (left) and signalling model (right) with putative *Coptis* orthologues annotated; bottom, heatmap of key gene expression in rhizomes across species.(H) Subcellular localization of *BAK1*–GFP and *BRI1*–GFP in epidermal cells (YFP channel, bright field image and overlay image).(I) Bimolecular fluorescence complementation (BiFC) assays testing *BRI1*–*BAK1* interactions (with controls).(J) Yeast two-hybrid assays of protein interactions between *BRI1* and *BAK1*.

To further investigate whether BRs are involved in regulating rhizome development in *C. chinensis*, we quantified endogenous BR levels in the rhizomes of four species. The results revealed a significant increase in the BR content in *C. chinensis* (Fig. 5C–5E and Supplementary Table15). Comparative transcriptomic analysis revealed that several key enzymes in the BR biosynthesis pathway, such as *CPD* and *BR6ox1*, were highly expressed in *C. chinensis* (Fig. 5F, heatmap and Supplementary Table16-19). In the BR signalling pathway, we also observed significantly elevated transcript levels of the core receptor complex components *BAK1* and *BRI1*, as well as the downstream transcriptional regulators *BZR1* and *BES1*, suggesting enhanced BR perception and signal transduction in *C. chinensis*.

To explore whether the molecular mechanism of BR-mediated rhizome development is conserved in *C. chinensis*, we performed subcellular localization experiments, which revealed that both *BRI1* and *BAK1* are localized to the plasma membrane. Subsequent bimolecular fluorescence complementation and yeast two-hybrid assays supported a physical interaction between *BRI1* and *BAK1* at the plasma membrane in response to BR induction, which may influence the regulation of downstream targets associated with rhizome expansion (Fig. 5H–5J).

Together, these results suggest that the coordinated activation of BR biosynthesis and signalling pathways in *C. chinensis* may enhance cellular expansion and branching and thus contribute to rhizome development and improving biomass accumulation and medicinal yield.

### Evolution and expression regulation of key genes involved in Coptis alkaloid biosynthesis

The core pharmacologically active compounds of *C. chinensis* are protoberberine-type alkaloids specifically enriched in the rhizome, mainly including berberine, coptisine, palmatine, and jatrorrhizine. Previous studies have preliminarily identified nine key enzyme families involved in their biosynthesis. To investigate the evolutionary expansion patterns of key enzyme gene families related to alkaloid biosynthesis in *Coptis*^74-75^, we selected four *Coptis* species (*C. chinensis*, *C. deltoidea*, *C. omeiensis*, and *C. teeta*), together with medicinal plants known to produce isoquinoline alkaloids (*Papaver somniferum* and *Phellodendron amurense*), as well as outgroup species (*Zanthoxylum bungeanum*, *Citrus maxima*, and *Scutellaria baicalensis*), for homolog identification of the nine key enzyme genes. The results showed that most key gene families involved in benzylisoquinoline alkaloid (BIA) biosynthesis, including (*S*)-norcoclaurine synthase (NCS), the *O*-methyltransferase family (6*O*MT, 4′*O*MT, S*O*MT), berberine bridge enzyme (BBE), and the CYP719A subfamily of cytochrome P450, underwent significant lineage-specific expansion in *Coptis* species (Figure 6A). This expansion pattern may provide an important genetic basis for the efficient biosynthesis of protoberberine alkaloids in *Coptis*.

**Fig. 6.**
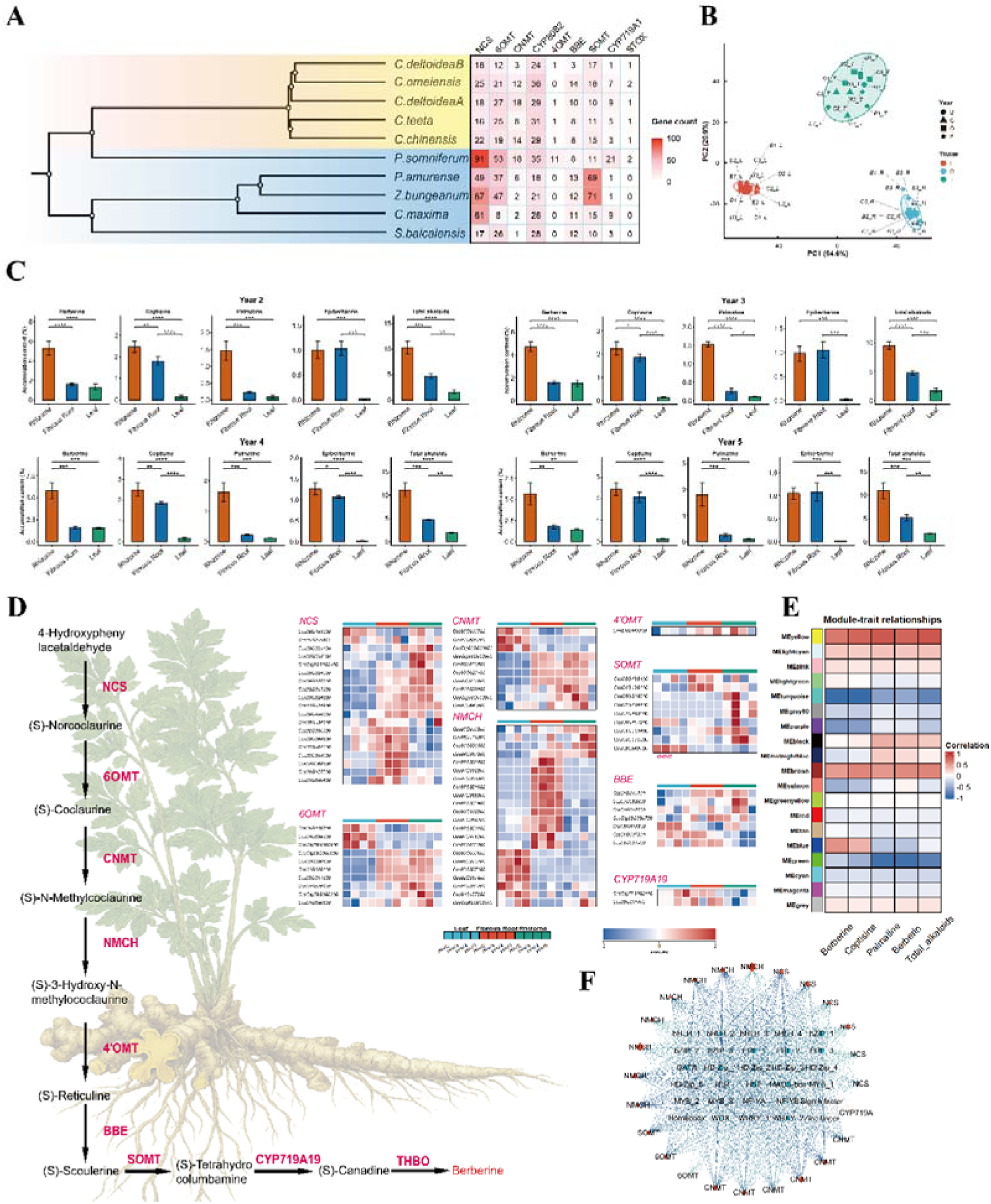
Analysis of alkaloid biosynthesis in *Coptis*. (A) Phylogenetic relationships and gene copy number distribution of nine key alkaloid biosynthetic enzyme families in *Coptis* and related species.(B) Principal component analysis (PCA) of transcriptomes from different tissues (leaf, fibrous root, and rhizome) of 2–5-year-old *Coptis* plants.(C) Contents of berberine, coptisine, palmatine, epiberberine, and total alkaloids in different tissues of 2–5-year-old *Coptis* plants.(D) Schematic diagram of the protoberberine alkaloid biosynthetic pathway in *Coptis* and heatmaps showing the expression of key structural enzyme genes in different tissues.(E) Heatmap of correlations between WGCNA co-expression modules and alkaloid content traits.(F) Putative co-expression network between key alkaloid biosynthetic structural genes and candidate regulatory factors.

We examined the spatial and temporal expression patterns of alkaloid biosynthesis-related genes by performing transcriptome sequencing on three major tissues from 2–5-year-old *Coptis* plants, including rhizome, fibrous root, and leaf. Principal component analysis showed that tissue type was the main factor driving gene expression variation. Leaf (L), fibrous root (R), and rhizome (T) samples formed clearly separated clusters, and the first two principal components (PC1 and PC2) together explained 85.5% of the total expression variation (Figure 6B). Samples from the same tissue but different growth years were closely grouped, indicating that the effect of growth year on gene expression was much smaller than that of tissue type. At the same time, the contents of berberine, coptisine, palmatine, and epiberberine were quantitatively measured in the corresponding samples. The results showed that both the contents of the four major alkaloids and the total alkaloid content were significantly higher in the rhizome than in the fibrous root and leaf, indicating that the rhizome is the main medicinal tissue for alkaloid accumulation. The fibrous root still contained certain levels of berberine and coptisine, whereas alkaloid contents in the leaf were extremely low(Figure 6C).

Based on the identified functional genes involved in alkaloid biosynthesis, we completely reconstructed the biosynthetic pathway of protoberberine-type alkaloids in *Coptis* and generated the corresponding expression heatmap. The results showed that different enzyme genes displayed clear differential expression across tissues and developmental stages (Figure 6D). Overall, upstream genes such as NCS and 6*O*MT were expressed in all three tissues, but their expression levels were significantly higher in the rhizome. In contrast, downstream modification enzymes, such as BBE and CYP719A19, showed much stronger tissue specificity and were mainly highly expressed in the rhizome. This expression pattern of “broad upstream expression but enhanced downstream tissue-specific expression” suggests a hierarchical transcriptional regulation of alkaloid biosynthesis.

To further explore the potential molecular regulatory mechanism underlying the biosynthesis of core alkaloids such as berberine, we performed weighted gene co-expression network analysis and focused on identifying key modules highly associated with alkaloid content traits. Using strict thresholds for gene significance and module membership, a total of 774 hub genes were identified, including 24 key structural genes involved in alkaloid biosynthesis and 30 transcription factors (Figure 6E). Based on co-expression relationships, we constructed a putative regulatory network linking transcription factors with alkaloid biosynthetic genes. In this network, transcription factors belonging to the MYB, bHLH, and ERF families exhibited highly connected hub-node characteristics and were strongly co-expressed with multiple key genes involved in alkaloid biosynthesis (Figure 6F). These transcription factors can therefore be considered candidate regulatory factors associated with protoberberine alkaloid biosynthesis in *Coptis*. Taken together, these results suggest that protoberberine alkaloid biosynthesis is associated with a coordinated candidate regulatory network involving multiple transcription-factor families; however, these regulators remain to be experimentally validated.

### Construction of the *Coptis* multiomics database

To fill the gap in platforms for the online integration and analysis of high-throughput sequencing datasets for *Coptis* species, we constructed the first *Coptis* Multiomics Database (https://Coptis-211.bioinfotoolkits.net:8443/). This database integrates high-throughput sequencing data, gene annotation information, expression profile analysis results, and related metabolic pathway information for four *Coptis* species (*C. chinensis, C. deltoidea, C. omeiensis* and *C. teeta*), providing data support for genetic research, exploration of medicinal ingredient biosynthesis mechanisms, and molecular breeding of *Coptis* species.

## Discussion

The availability of high-quality reference genomes is fundamental for understanding the evolutionary history and trait diversification of medicinal plants. Although genome assemblies of *C. chinensis* and *C. teeta* have previously been reported^9,10^, limitations in assembly continuity and completeness have restricted chromosome-scale comparative analyses. By integrating PacBio HiFi sequencing, Hi-C scaffolding, and comparative genomics, we generated four chromosome-level genomes representing the major evolutionary lineages of *Coptis*. Compared with previously published assemblies, these genomes exhibit substantially improved continuity and completeness, providing a robust framework for investigating genome evolution, adaptive diversification, and the genetic basis of medicinally important traits. Moreover, the simultaneous comparison of four closely related species enables reconstruction of evolutionary processes that could not be resolved using single-species genomic resources alone.

Our phylogenomic analyses suggest that the evolutionary history of *Coptis* has been closely associated with major geological and climatic changes in East Asia. Divergence-time estimation indicated that the split between *Coptis* and other Ranunculaceae lineages occurred approximately 73.9 Mya, coinciding with the early stages of the India-Asia collision and subsequent environmental changes across Eurasia^23-25^. Diversification within *Coptis* mainly occurred during the Miocene and late Miocene-Pliocene, overlapping with major phases of Tibetan Plateau uplift, Asian monsoon intensification, and Neogene cooling^14,15,23^. Notably, each major divergence event was accompanied by extensive gene-family expansion and contraction, suggesting that genomic innovation may have facilitated adaptation to changing environments. However, the relationship between geological events and species diversification should be interpreted cautiously. In addition to geological and climatic processes, ecological specialization, demographic history, and biotic interactions may also have contributed to lineage divergence. Future population genomic and phylogeographic studies will be necessary to disentangle the relative importance of these factors.

Against this geological background, polyploidization and subsequent subgenome differentiation may represent an important genomic route by which *Coptis* lineages acquired evolutionary novelty. Triploidization provides an important source of genomic variation during plant evolution, yet different subgenomes may follow distinct evolutionary trajectories. In *C. deltoidea*, we observed pronounced expression dominance of the B subgenome together with contrasting patterns of chromosomal evolution between the two ancestral genomic components. Whereas the B subgenome largely retained its chromosomal organization, the A subgenome experienced multiple chromosome breakage and rearrangement events. Similar homoeolog expression bias and expression-level dominance have been documented in other allopolyploid crops, including hexaploid wheat and allopolyploid cotton^30,31^. In synthetic and natural allopolyploids, subgenome-biased expression can emerge rapidly after hybridization and has frequently been linked to asymmetric transposable-element landscapes, differential TE methylation, and gene-proximal epigenetic states^32,33^. Therefore, the B-subgenome expression dominance observed in *C. deltoidea* is unlikely to be an isolated phenomenon; rather, it may represent a convergent outcome of allopolyploid genome stabilization. Nevertheless, the AAB triploid constitution and the marked structural conservation of the B subgenome distinguish *C. deltoidea* from classical allotetraploid systems, suggesting that the strength and direction of dominance may also reflect lineage-specific dosage balance, chromosomal stability, and regulatory-network rewiring. Future methylome profiling, chromatin-accessibility assays, and subgenome-specific TE annotation will be required to determine whether B-subgenome dominance in *C. deltoidea* is primarily driven by the same epigenetic and TE-related mechanisms observed in other polyploids or by a distinctive triploid-specific regulatory architecture.

In addition to genome-wide expression asymmetry, immune-gene evolution provides another axis through which *Coptis* may have adapted to heterogeneous mountain environments. NLR genes are central components of plant innate immunity and play critical roles in pathogen recognition and defense responses^18-20^. Comparative analyses revealed substantial expansion of NLR gene families in *C. omeiensis* and the A subgenome of *C. deltoidea*, particularly through large-scale duplication events. An intriguing feature of all four *Coptis* genomes is the absence of complete TNL-type NLR genes under our annotation pipeline. The angiosperm NLR atlas reported large copy-number variation among closely related species and showed that NLR reduction is associated with ecological specialization and deletion of downstream immune-signalling components^20^. More recent synteny-informed analyses further indicate that independent TNL losses have occurred repeatedly during angiosperm evolution and are often coupled with changes in EDS1-family and helper-NLR signalling modules^26^. In this context, the lack of complete TNL genes in *Coptis* should be interpreted as a possible lineage-specific reorganization of immune signalling rather than a simple stochastic loss. Because *Coptis* species are perennial herbs with underground rhizomes continuously exposed to soil-borne microorganisms, the expansion of non-TNL NLR classes may represent compensatory diversification of immune recognition capacity. Nevertheless, the adaptive significance of NLR expansion remains largely speculative, and direct functional evidence linking NLR copy-number variation to pathogen resistance or ecological specialization is still required.

The above evolutionary processes also connect to economically important morphological divergence. Among the four investigated species, *C. chinensis* is unique in developing a highly branched and enlarged rhizome, which contributes substantially to its medicinal yield and economic value. We found that *C. chinensis* accumulated significantly higher levels of brassinosteroids (BRs) than the other three species and exhibited elevated expression of the BR biosynthetic gene BR6ox1. Furthermore, the interaction between BRI1 and BAK1 suggests that enhanced BR signaling may promote downstream pathways involved in rhizome development^16,17,27-29^. These findings support a role for BR biosynthesis and signaling in the distinctive rhizome architecture of *C. chinensis*, although additional functional studies will be required to establish causality and determine how BR signaling interacts with other developmental pathways.

In addition to biomass-related traits, medicinal quality in *Coptis* depends largely on specialized metabolism. Protoberberine alkaloids are the principal bioactive compounds responsible for the medicinal properties of *Coptis*^1,3,9^. Comparative analyses revealed extensive expansion of several key enzyme families involved in alkaloid biosynthesis^74,75^, together with pronounced rhizome-specific expression patterns. These results suggest that both gene-family expansion and transcriptional specialization have contributed to the efficient accumulation of alkaloids during the evolutionary history of the genus. In addition, co-expression analyses identified multiple transcription-factor families, including MYB, bHLH, and ERF, as candidate regulatory factors associated with alkaloid biosynthesis, suggesting that metabolite production may be controlled by a multilayered regulatory network. However, these regulatory relationships remain largely inferred from co-expression patterns and require experimental validation.

Despite the comprehensive genomic and multi-omics analyses presented here, several limitations should be acknowledged. The associations between geological events and species diversification remain largely correlative, the molecular basis of subgenome dominance cannot yet be resolved in the absence of epigenomic and TE-distribution data, and the functional significance of NLR expansion and candidate regulatory factors remains to be experimentally verified. Future studies integrating population genomics, epigenomics, functional genetics, and genome editing will provide deeper insights into the mechanisms underlying adaptive evolution, medicinal trait formation, and species diversification in *Coptis*.

## Methods

### Plant materials and sequencing

The leaves, stems, roots, fruits, and flowers of *C. chinensis, C. deltoidea, C. omeiensis* and *C. teeta* were collected from Shizhu *Coptis* Germplasm Resource Nursery. These samples were frozen in liquid N_2_ and preserved at -80 °C for use in subsequent DNA and RNA extractions. A modified cetyltrimethylammonium bromide (CTAB) technique was utilized to extract high-quality genomic DNA^34^. A commercial RNA extraction kit (TIANGEN, Beijing, China) was used to extract RNA.

A short-read (150 bp) library was subsequently produced and sequenced on a DNBSEQ platform (BGI, MGISEQ-2000). Circular consensus sequencing (CCS) libraries were produced and sequenced on a PacBio Sequel II platform (Pacific Biosciences, CA, USA). High-throughput Chromosome Conformation Capture (Hi-C) fragment libraries were produced using a range of insert sizes (300–700 bp)^35^ and sequenced on DNBSEQ PE150. Finally, the extracted RNA was also sequenced using a DNBSEQ PE150 (BGI), and the resulting data were used for library prediction.

### Genome assembly and evaluation

The PacBio HiFi sequencing data were assembled using Hifiasm^36^ to generate the initial contigs. The quality-filtered Hi-C data were subsequently aligned to the draft genome assembly with BWA^37^, and the alignment results were subsequently analysed using Juicer^38^ to construct a three-dimensional interaction map of the genome. Based on this interaction information, 3D-DNA^39^ was employed to break misjoined contigs and anchor them onto pseudochromosomes. Thereafter, JuiceBox^40^ was used to visualize the interaction relationships within and between pseudochromosomes, allowing manual correction of potential misassemblies. The corrected interaction information was then reprocessed with 3D-DNA, ultimately yielding a high-quality chromosome-level genome assembly. The quality of the assembly was assessed using BUSCO^41^ to evaluate gene completeness, the LTR Assembly Index (LAI)^42^ to assess continuity and completeness in repetitive regions, and CGEMA^43^ to comprehensively evaluate the overall quality and accuracy of the assembled genome.

### Genome annotation

Repetitive sequences in the genome were annotated using a combination of de novo and homology-based approaches. In the de novo prediction, RepeatModeler^44^ and LTR-FINDER^45^ were used to construct a repeat sequence library, which was subsequently subjected to RepeatMasker^46^ to predict repetitive elements. Tandem Repeat Finder^47^ was further employed to detect and identify tandem repeats in the genome. For the homology-based prediction, RepeatMasker was used to align the genome sequences against the Repbase database, thereby identifying homologous repetitive elements. Finally, the results from both approaches were integrated to produce a comprehensive and high-quality annotation of repetitive genomic sequences.

Gene structure annotation was performed by combining transcriptome-based prediction, homology-based prediction, and ab initio prediction. For ab initio prediction, Augustus^48^ and SNAP^49^ were used to predict potential gene models directly from the genome sequence. For homology-based prediction, GeMoMa^50^ was used to align the genome sequence with protein sequences from *Aquilegia coerulea*, *Macleaya cordata*, and *Corydalis edulis Maxim* to infer putative coding genes. For transcriptome-based prediction, Trinity^51^ was used to perform de novo assembly of RNA-seq data to obtain transcript sequences, which were further aligned to the genome and refined using PASA^52^ to improve gene structure annotation. Finally, the results from the three approaches were integrated using EvidenceModeler^52^, which weighted and combined evidence from different sources to generate a comprehensive and high-quality gene structure annotation.

Gene functional annotation was primarily performed by aligning protein-coding genes against various functional databases to predict their potential functions. In this study, the functional databases used included Nr, SwissProt, KEGG, COG, TrEMBL, and InterPro.

### Transcriptome analyses

FastQC^53^ was used to assess the quality of the raw RNA sequencing data, and Trimmomatic^54^ was employed to remove low-quality reads and adapter sequences. High-quality clean reads were subsequently aligned to the reference genomes of the four *Coptis* species using STAR^55^. The alignment results were converted into sorted BAM format files with SAMtools^56^. Afterwards, StringTie^57^ was used to assemble the transcripts and quantify the gene expression levels from the alignment data. Finally, the read counts were normalized to both the gene length and the total number of mapped reads in each sample to calculate TPM (transcripts per million) values, which were used to represent gene expression levels.

### Gene family clustering analysis

We collected the genome and annotation data of *Aquilegia vulgaris*, *Epimedium sagittatum*, *Oryza sativa*, *Papaver somniferum*, and *Phellodendron amurense* from the NCBI database, together with the four *Coptis* species sequenced in this study. Redundant transcript isoforms were removed using Gffread^58^, and only the longest transcript was retained for each gene. Gene family clustering was performed with OrthoFinder^59^ using the DIAMOND^60^ alignment mode for sequence similarity searches, while all other parameters were set to default values.

### Phylogenetic analysis and divergence time estimation

Single-copy orthologous genes shared by nine species were identified through clustering. These genes were aligned using MAFFT^61^, and the resulting alignments were concatenated to construct a supergene sequence. A phylogenetic tree was then inferred with RAxML based on the maximum likelihood method, with 1000 bootstrap replicates. On this basis, species divergence times were estimated using MCMCTree^62^ in the PAML package and calibrated with fossil-based time points obtained from the TimeTree^63^ online database. Finally, the phylogenetic tree was visualized and refined using the Itol^64^ platform.

### Gene family expansion and contraction analysis

CAFE^65^ software was used to estimate the ancestral gene family sizes for each branch based on a birth–death model, thereby predicting the expansion and contraction of gene families relative to their ancestors. KEGG and GO enrichment analyses were then performed for the expanded and contracted gene families, with P < 0.05 considered to indicate statistical significance.

### Positive selection analysis

Based on single-copy genes, the codeml program in the PAML^66^ package was used to calculate the Ka/Ks ratio (nonsynonymous to synonymous substitution rate) under both the branch model and the branch-site model. Subsequently, *p* values were computed based on chi-square statistics, and genes with *p* < 0.05 were considered to have undergone strong positive selection. The positively selected genes identified were then subjected to GO and KEGG enrichment analyses.

### Ancestral karyotype reconstruction

To trace the karyotype evolution of the four *Coptis* species, we first used MCScan^67^ to compare the *Coptis* genomes with the *Tetracentron sinense* genome and identified 1:1 homologous sequences. Next, DRIMM-Synteny^68^ was employed to filter and extract conserved syntenic blocks. Finally, the IAGS program^69^ (based on the GMP model) was applied to reconstruct the ancestral genome structures at key evolutionary nodes.

### Subgenome phasing and expression bias classification

Given that *C. deltoidea* has an AAB triploid genotype, we first used SubPhaser^70^ to phase its genome based on orthologous genes and subgenome-specific k-mers, thereby dividing it into two subgenomes: *C. deltoidea A* and *C. deltoidea B*. We then counted the number of homologous genes in each subgenome and calculated the average TPM values of homologous gene pairs across different tissues using transcriptome data to evaluate expression differences. Homologous gene pairs exhibiting expression bias were defined as genes whose expression was biased, indicating that the expression of a gene from one subgenome was significantly greater than that of its corresponding homologue in the other subgenome. According to the degree of bias, these gene pairs were classified into five categories: (i) A_sig_up: A-subgenome significantly biased expression gene pairs (log ^(TPMA+1)^/log ^(TPMB+1)^ ≥ 2); (ii) B_sig_up: B-subgenome significantly biased expression gene pairs (log ^(TPMA+1)^/log ^(TPMB+1)^ ≤ -2); (iii) A_subsig_up: A-subgenome moderately biased expression gene pairs (1 ≤ log ^(TPMA+1^)/log ^(TPMB+1)^ < 2); (iv) B_subsig_up: B-subgenome moderately biased expression gene pairs (-2 < log ^(TPMA+1)^/log ^(TPMB+1)^ ≤ -1); and (v) no diff: unbiased gene pairs (0 ≤ |log (TPMA+1)/log ^(TPMB+1)^| < 1). We then examined the distribution of these categories across different tissues, including leaves, roots, and stems. Finally, homologous gene pairs that exhibited significant expression differences across all the tissues were defined as dominant gene pairs, and GO enrichment analysis was performed separately for the dominant genes in the A and B subgenomes.

### Resistance gene identification and duplication analysis

RGAugury^71^ was used to identify resistance genes in the genome and classify them into RNL, CNL, NL, CN, and N types. For *C. omeiensis*, the identified NLR protein sequences were aligned with MAFFT, and a maximum-likelihood phylogenetic tree was constructed using IQ-TREE^72^. The expression levels of NLR genes across different tissues were extracted from gene expression files, and chromosomal coordinates were obtained from genome annotations for localization. The phylogenetic tree, expression patterns, and chromosomal distributions were then integrated and visualized in iTOL. In addition, DupGen_finder^73^ was used to classify resistance gene duplications into six categories: WGD (whole-genome duplication), TD (tandem duplication, two adjacent duplicated genes), PD (proximal duplication, duplicated genes separated by fewer than 10 genes), TRD (transposed duplication, duplicated genes composed of ancestral and new loci), DSD (dispersed duplication, neither adjacent nor collinear), and SL (singleton genes). All duplicated genes in *C. omeiensis* were further subjected to GO and KEGG enrichment analyses, with p < 0.05 as the significance threshold.

### Yeast two-hybrid, BiFC, and subcellular localization assays

In the Y2H assay, the Clontech Matchmaker™ Gold system was used to clone different domains of *BAK1* and *BRI1* (kinase domain and extracellular domain) into pGBKT7 (BD vector) and pGADT7 (AD vector), respectively, and cotransform them into the AH109 yeast strain. Cotransformants were selected on DDO (SD/-Leu/-Trp) plates, and interactions were tested on QDO (SD/-Ade/-His/-Leu/-Trp) plates. Only the combination of *BAK1* and *BRI1* kinase domains increased on QDO plates, indicating a specific interaction between the kinase domains.

In the BiFC assay, the full-length cDNA of *BAK1* was cloned and inserted into pSPYNE(R)173 (YN vector), and the full-length cDNA of *BRI1* was cloned and inserted into pSPYCE(MR) (YC vector). The constructs were subsequently introduced into *Nicotiana benthamiana* leaves via *Agrobacterium*-mediated transient expression. At 48–72 h post-infiltration, confocal microscopy revealed strong YFP fluorescence at the plasma membrane in the test group, while the negative control group showed no signal, further confirming the in vivo interaction between *BAK1* and BRI1.

In the subcellular localization assay, the full-length CDSs of *BAK1* and *BRI1* were cloned and inserted into the pCAMBIA1300-35S-GFP vector and transiently expressed in *N. benthamiana* leaves. Confocal imaging revealed that both *BAK1* and *BRI1* localized to the plasma membrane, confirming their subcellular localization.

### Phenotypic measurement and statistical analysis

The rhizome weight, optical density (OD), and endogenous BR content of the four *Coptis* species were measured, with three independent biological replicates set for each parameter. The experimental data are expressed as the mean ± standard deviation (SD) and were visualized using bar charts.

### Identification of alkaloid biosynthetic genes

According to published studies ^74-75^, sequences of key enzyme genes involved in the berberine biosynthetic pathway in *Coptis* were collected as reference sequences. Homologous genes were then searched in nine species using BLASTN, with an E-value cutoff of 1e-5. Candidate sequences with similarity higher than 80% were retained and further verified using NCBI BLASTN. Based on this screening and validation process, homologous genes related to alkaloid biosynthesis were finally identified in each species.

### Principal component analysis

TPM values were extracted from the gene expression data to construct an expression matrix. To reduce the effect of highly expressed genes on overall variation, TPM values were transformed using log ^(x+1)^. The transformed data were then standardized by Z-score normalization. Based on the normalized expression matrix, PCA was performed using the prcomp function in R for linear dimensionality reduction and extraction of the major sources of variation among samples. The first two principal components (PC1 and PC2) were visualized using ggplot2 to assess the overall transcriptomic differences among tissues and growth years

### Weighted gene co-expression network analysis

Weighted gene co-expression network analysis (WGCNA) was performed using the WGCNA package^76^ in R. The goodSamplesGenes function was employed to filter low-quality genes and samples, followed by the pickSoftThreshold function to determine the optimal soft-thresholding power for scale-free network construction. Co-expression modules were identified using the blockwiseModules function with parameters: networkType = “signed”, TOMType = “signed”, minModuleSize = “30”, and mergeCutHeight = “0.25”. Module eigengenes were calculated, and Pearson correlation analysis was conducted to evaluate the relationships between modules and alkaloid traits. Hub genes were identified based on the thresholds of |MM| > 0.8 and |GS| > 0.5. Finally, gene pairs with weight values exceeding 0.2 in the target module were extracted to construct and visualize the regulatory network between transcription factors and alkaloid biosynthetic genes using Cytoscape^77^.

### HPLC analysis

Rhizome, fibrous root, and leaf samples of *Coptis* were dried to constant weight, ground into fine powder, and extracted by ultrasonic-assisted extraction. The extracts were centrifuged and filtered before HPLC analysis. Standard curves were established using reference standards, and the contents of berberine, coptisine, palmatine, epiberberine, and total alkaloids were calculated using the external standard method. Statistical differences were analyzed by one-way ANOVA, and bar plots were generated accordingly.

### Database implementation

The Coptis Multiomics Database (CMD) was constructed based on a modular web framework to ensure the efficient integration and analysis of multiomics data. The back end uses MySQL^78^ as the data storage and management system to support the stable storage and retrieval of multiple data sources, including genome assemblies, gene annotations, expression profiles, and metabolic pathways. Before being imported into the database, the data files were standardized and indexed to achieve rapid search and efficient access. For functional and comparative analyses, the platform integrates commonly used bioinformatics tools such as OrthoVenn3^79^ (for orthologous gene cluster identification), and GO enrichment analysis^80^ (for functional annotation and significance analysis). The frontend interface was designed using Vue (https://cn.vuejs.org/index.html) and Ant Design (https://ant.design/index-cn), providing good interactivity and a good user experience. Data and results visualization was achieved using ECharts^81^ and Canvas^82^. The platform has been optimized for modern browsers that support HTML5 (such as Google Chrome, Firefox, Safari, and Opera) to ensure efficient database interaction. The database is fully open and can be accessed without registration or login, providing researchers with a convenient environment for data acquisition and analysis.

## Data availability

The data that support the findings of this study are available in the supplementary material of this article. The sequencing data have been deposited on ngdc.cncb.ac.cn, The data are publicly accessible at: https://bigd.big.ac.cn/gsa/browse (Bio-project Accession ID: PRJCA046294).

## Acknowledgements

This work was financially supported by National Key R&D Program of China (2022YFD1201602); National Modern Agricultural Park Construction Project; Key project at central government level: The ability establishment of sustainable use for valuable Chinese medicine resources (2060302); Project for Transformation and demonstration of Scientific and Technological Achievements from Sichuan Province (24ZHSF0335); National Natural Science Foundation of China (82304837)

## Author contributions

Xiaoli Ye and Hang Ma conceived and designed the project. Mingjiang Huang performed most of the experiments, analyzed the data, and prepared the figures. Yingchun Zhang and Lei Pang conducted the yeast two-hybrid and BiFC assays. Mengguo Ran, Shenyong Chen, Yufeng Ruan, Wanqing Liu, Tingrui Peng, Tao Yu, Jian Qin, Difeng Tan, Xuefeng Yu, Xin Tang, Guanglou Nie, and Pingan Peng contributed to sample collection and growth measurements. Mengguo Ran, Tingrui Peng, and Pingan Peng also contributed to brassinosteroid (BR) quantification. Xing Zhao and Xuegang Li analyzed the BR biosynthetic pathway. Mingjiang Huang drafted the manuscript. Hang Ma and Xiaoli Ye supervised the project and revised and finalized the manuscript with input from all authors.

## Supplementary figures

**Supplementary Fig. 1.**
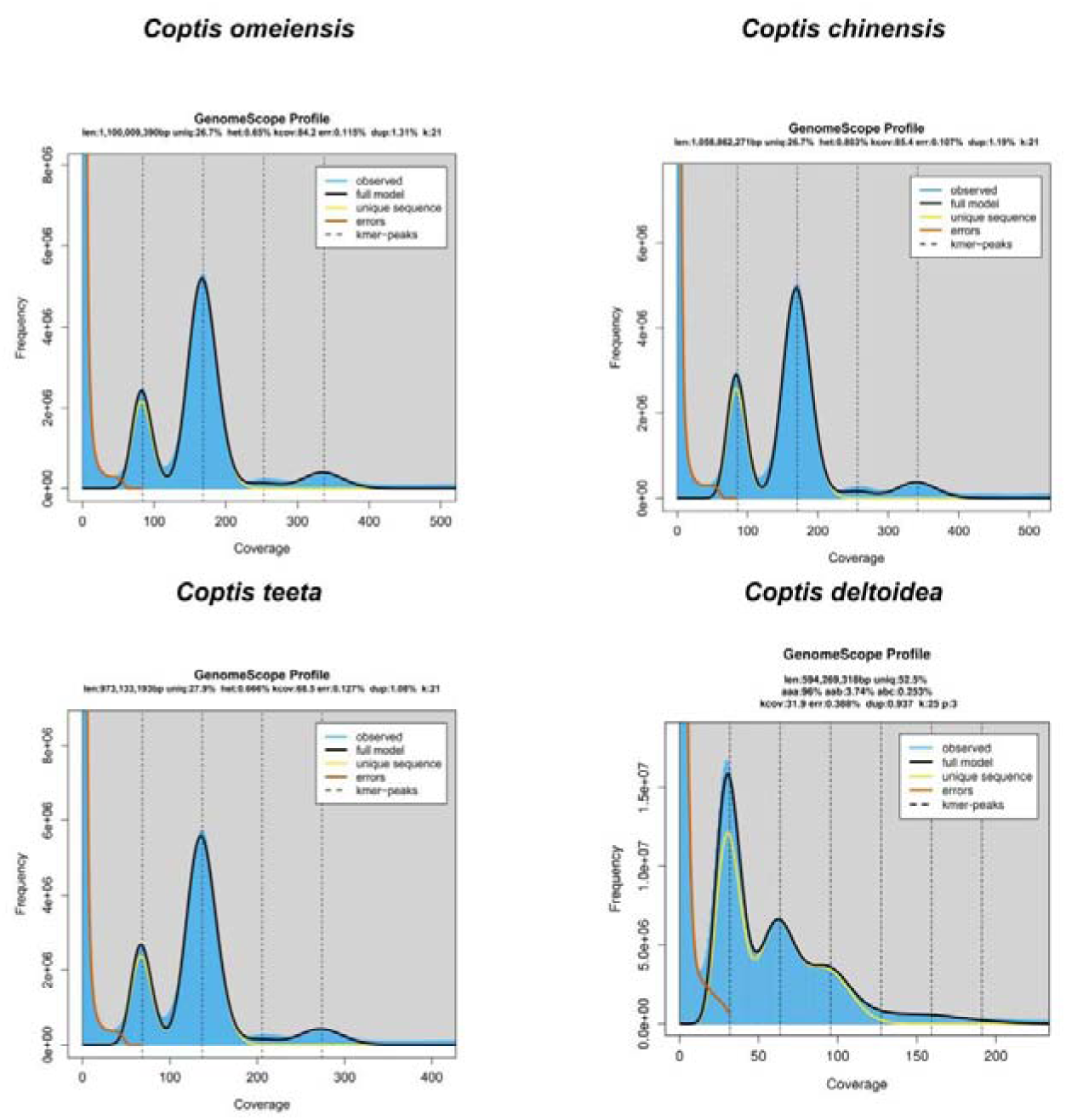
GenomeScope profiles of four *Coptis* species.

**Supplementary Fig. 2.**
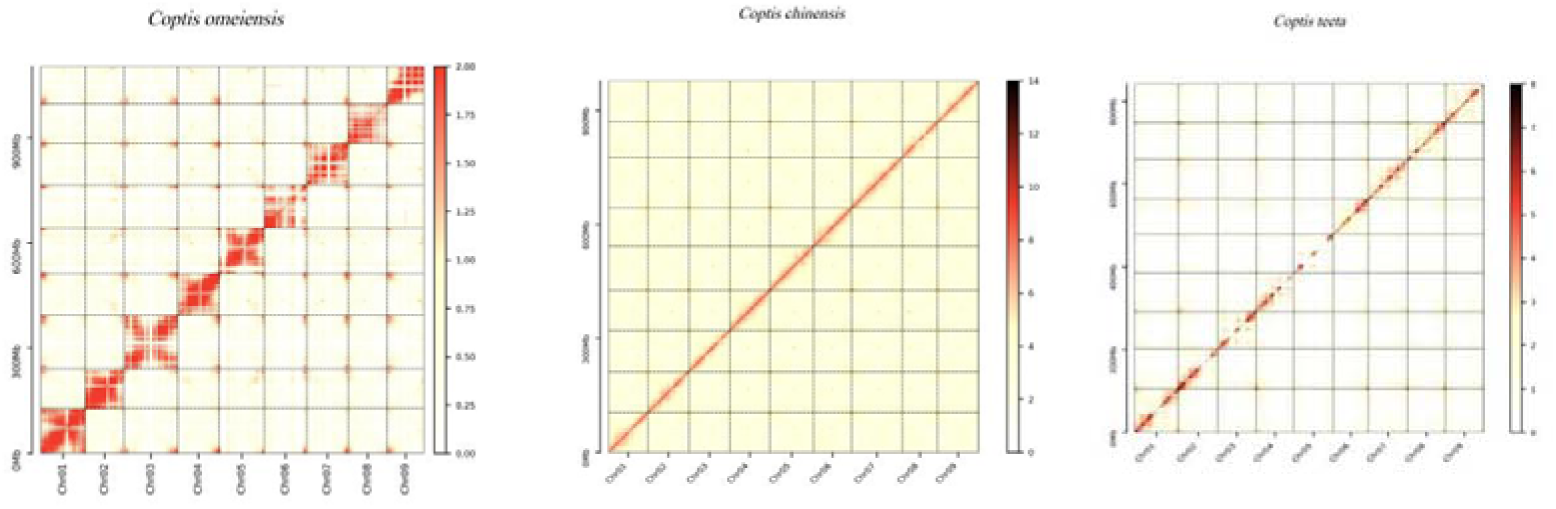
Hi-C contact heatmaps for the chromosome-level genome assemblies of three *Coptis specie*s.

**Supplementary Fig. 3.**
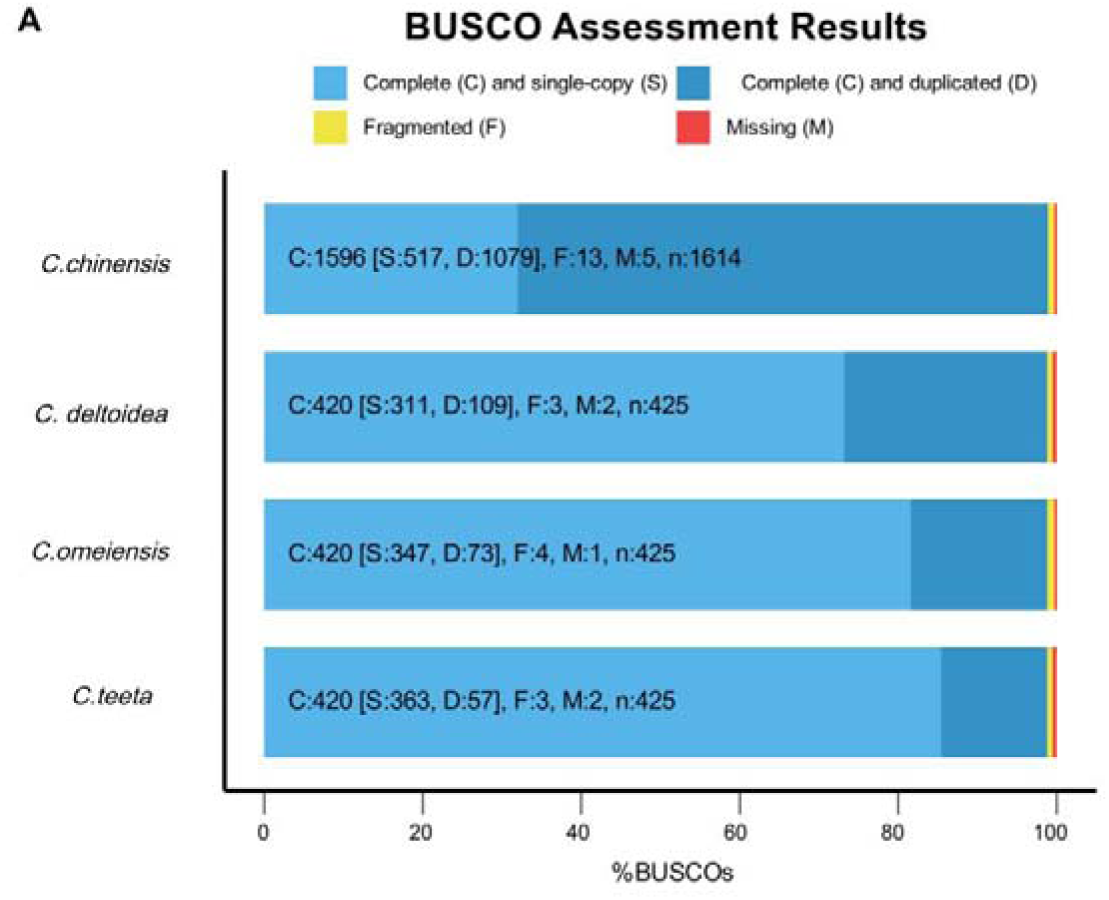
BUSCO assessment of the four Coptis genome assemblies.

**Supplementary Fig. 4.**
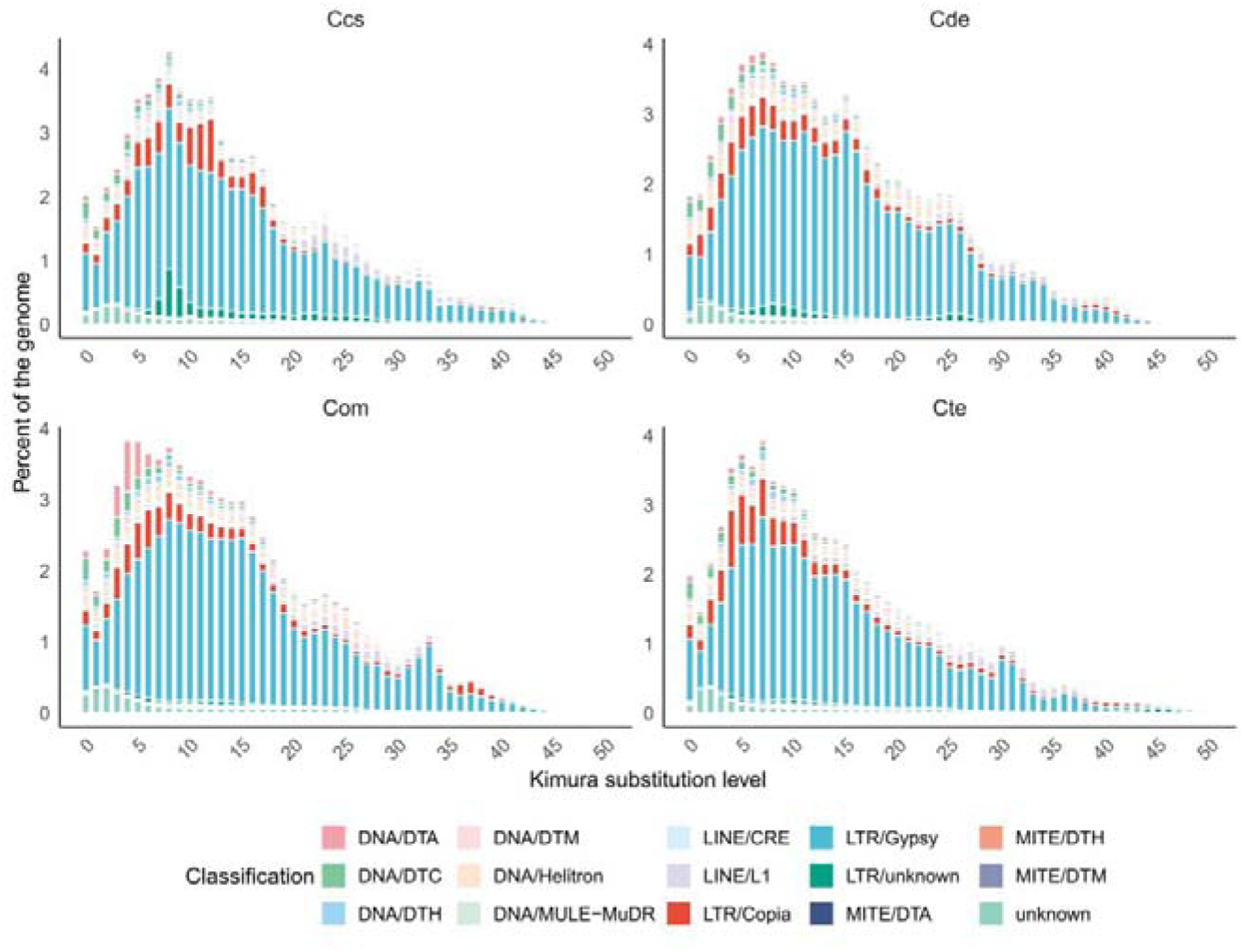
Kimura divergence distribution of transposable elements (TEs) in the four *Coptis* genomes.

**Supplementary Fig.5.**
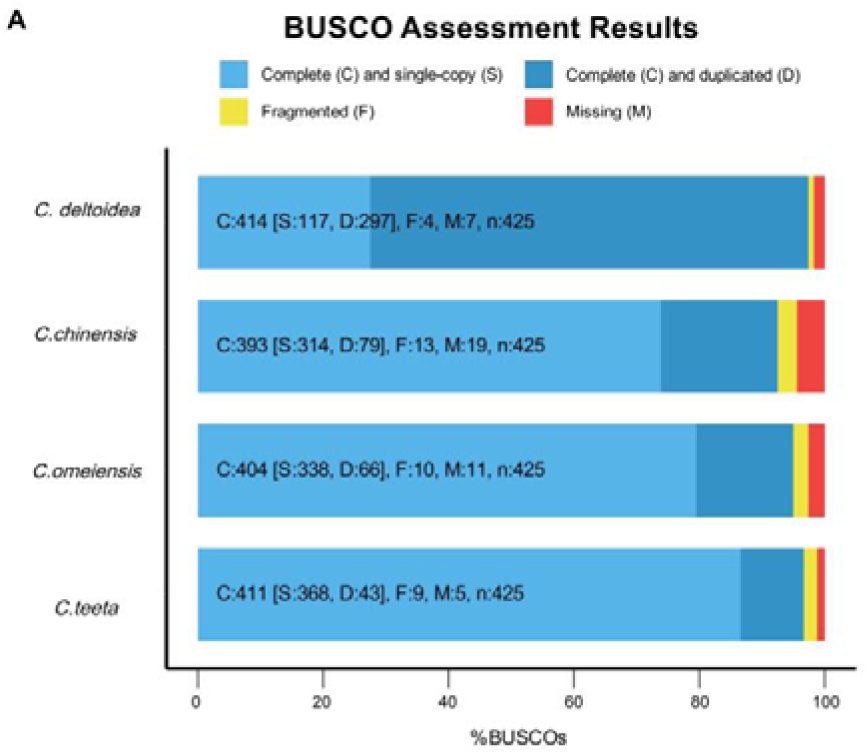
BUSCO assessment of predicted protein-coding genes in the four Coptis genomes.

**Supplementary Fig. 6.**
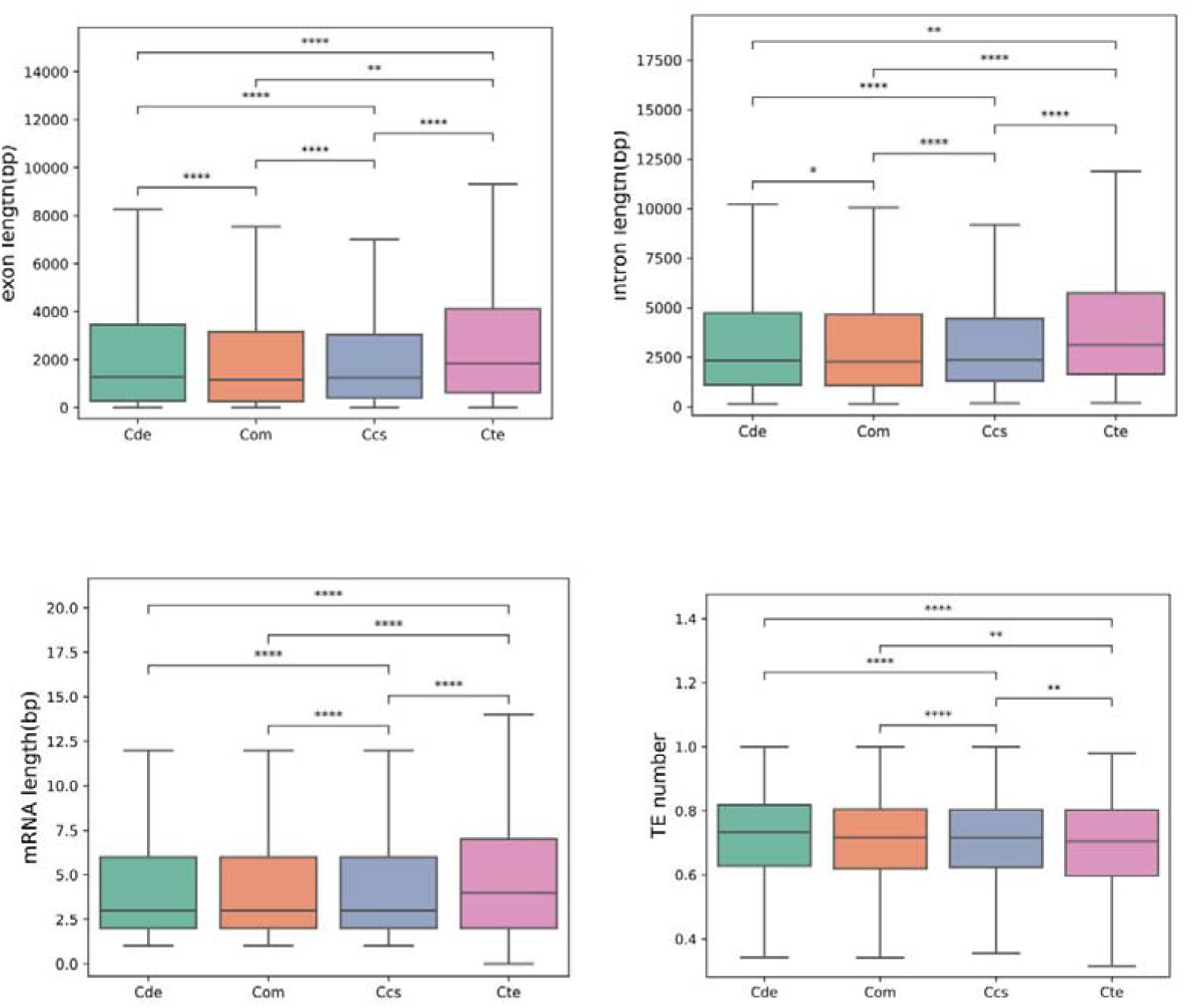
Statistical characterization of genomic features and repetitive elements in four *Coptis* species.

**Supplementary Fig. 7.**
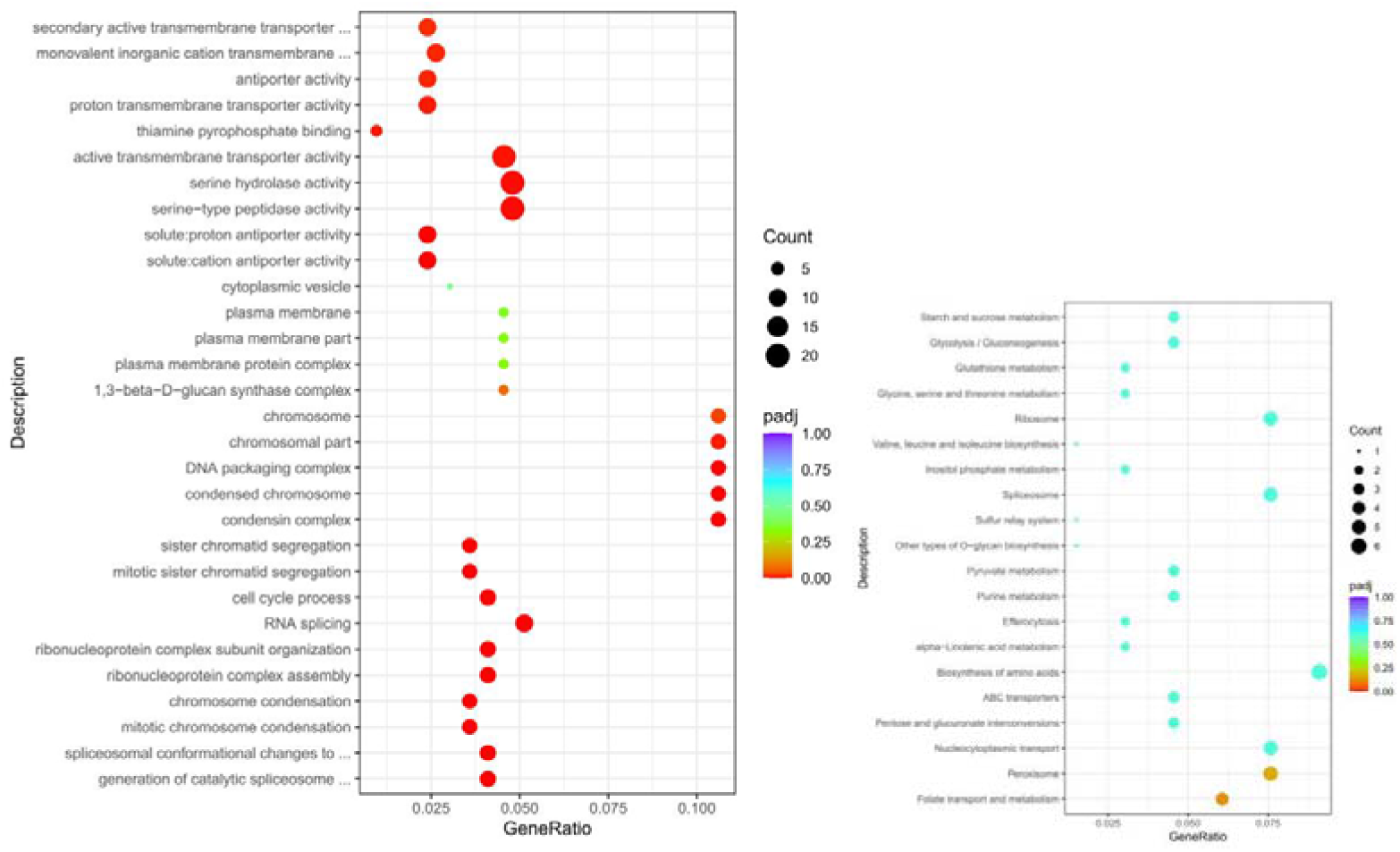
Functional enrichment analysis of unique gene families in *Coptis chinensis*.

**Supplementary Fig. 8.**
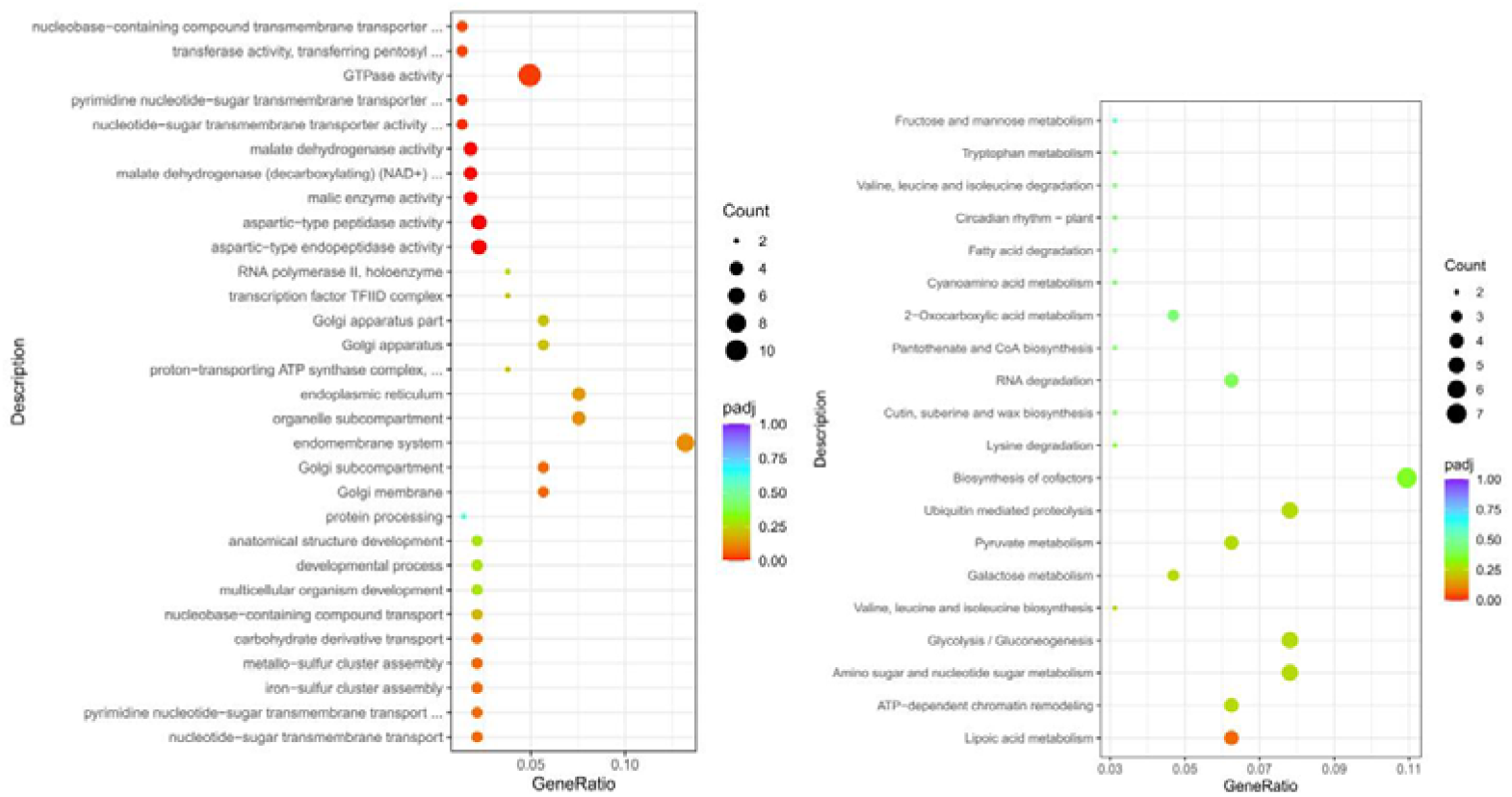
Functional enrichment analysis of unique gene families in ***Coptis deltoidea* subgenome A**.

**Supplementary Fig. 9.**
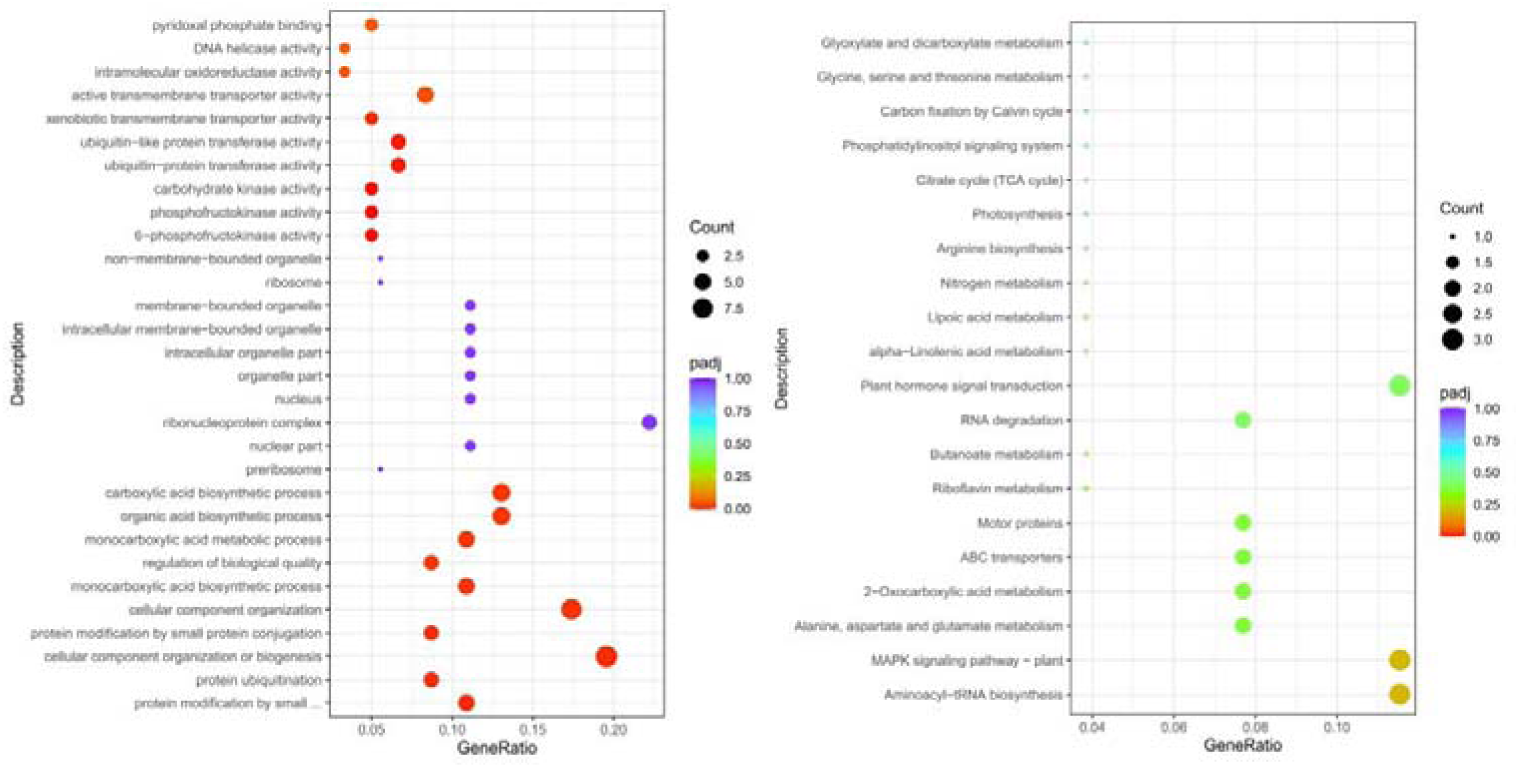
Functional enrichment analysis of unique gene families in Coptis deltoidea subgenome B.

**Supplementary Fig.10.**
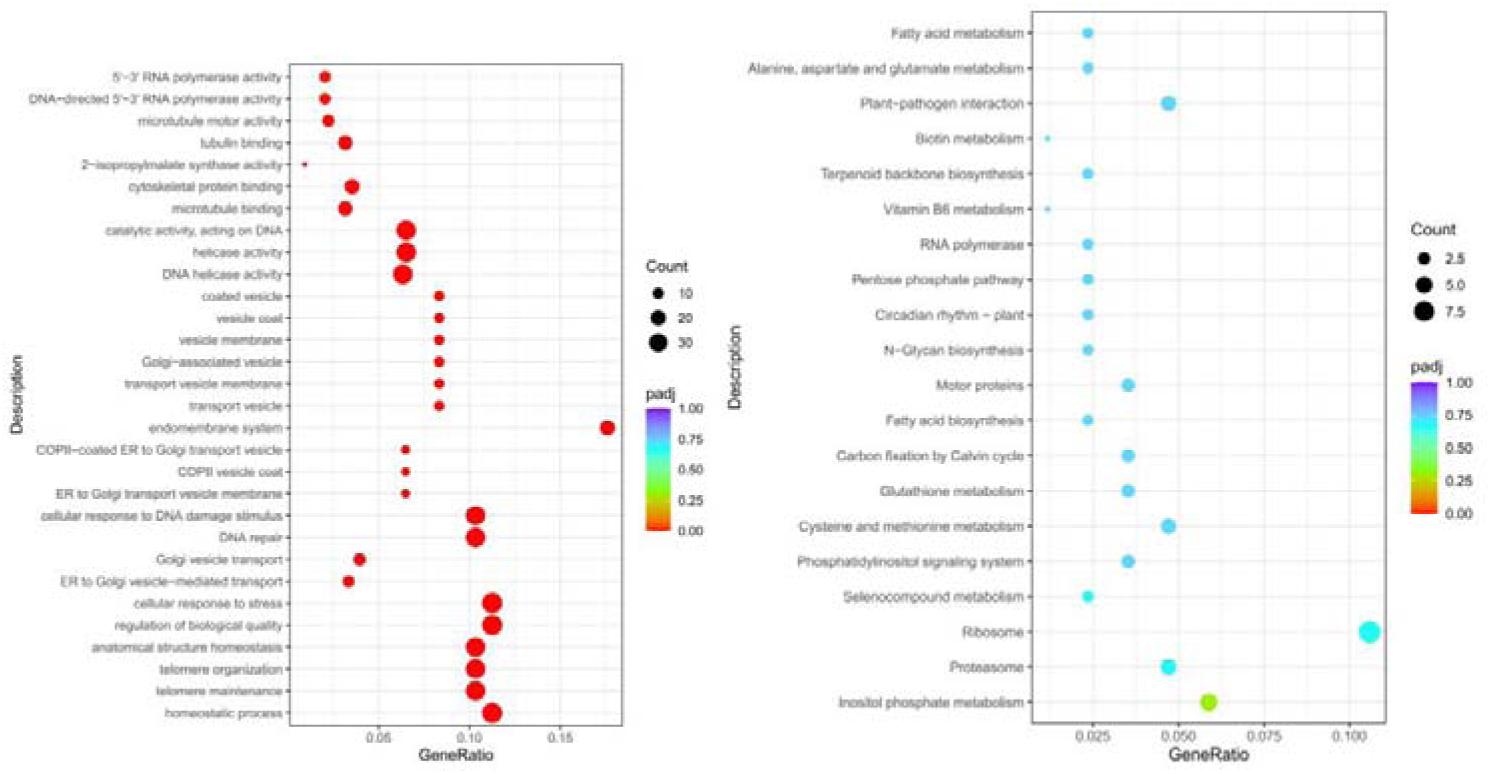
Functional enrichment analysis of unique gene families in *Coptis teeta*.

**Supplementary Fig.11.**
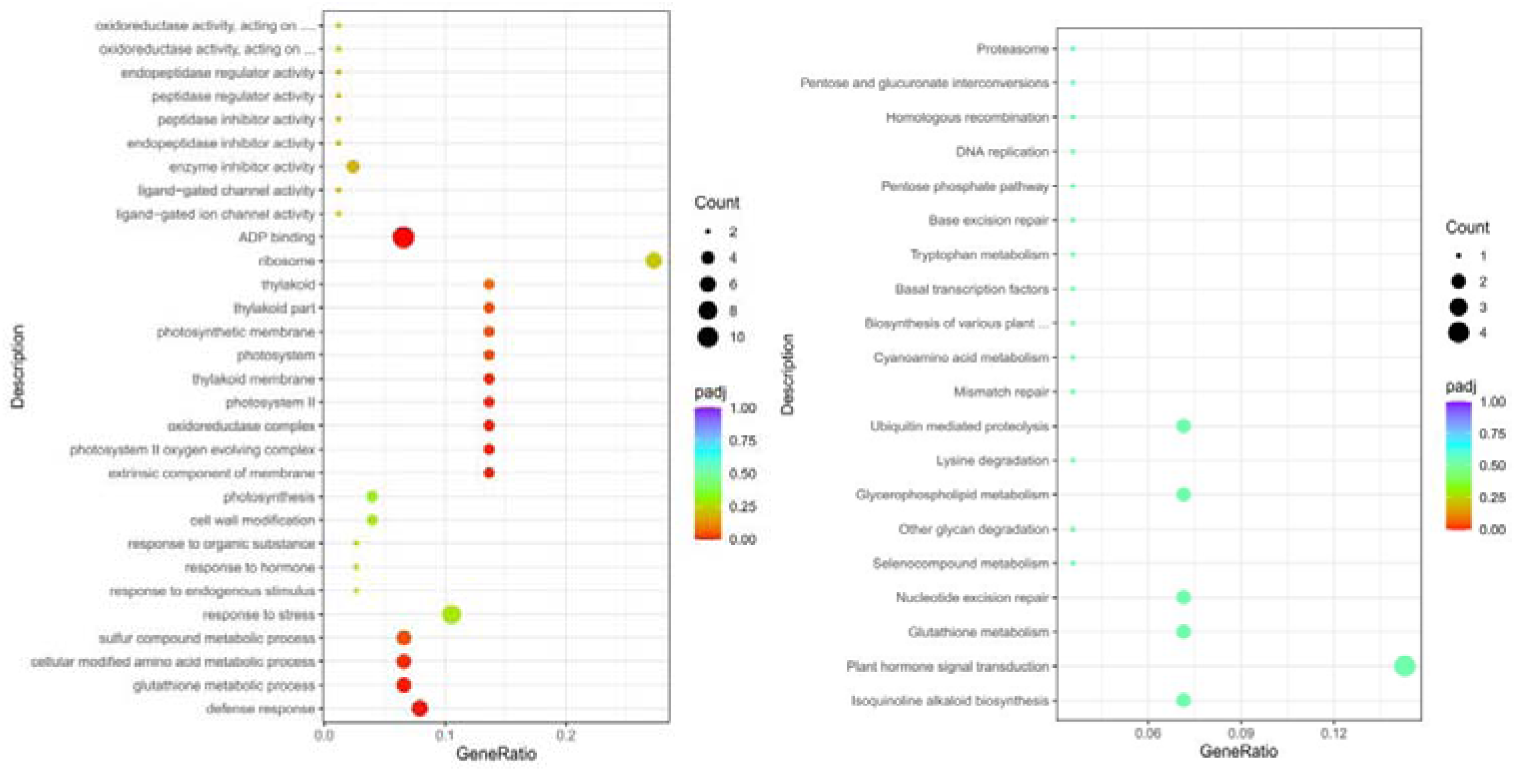
GO and KEGG enrichment analysis of significantly expanded gene families in *C. chinensis*.

**Supplementary Fig.12.**
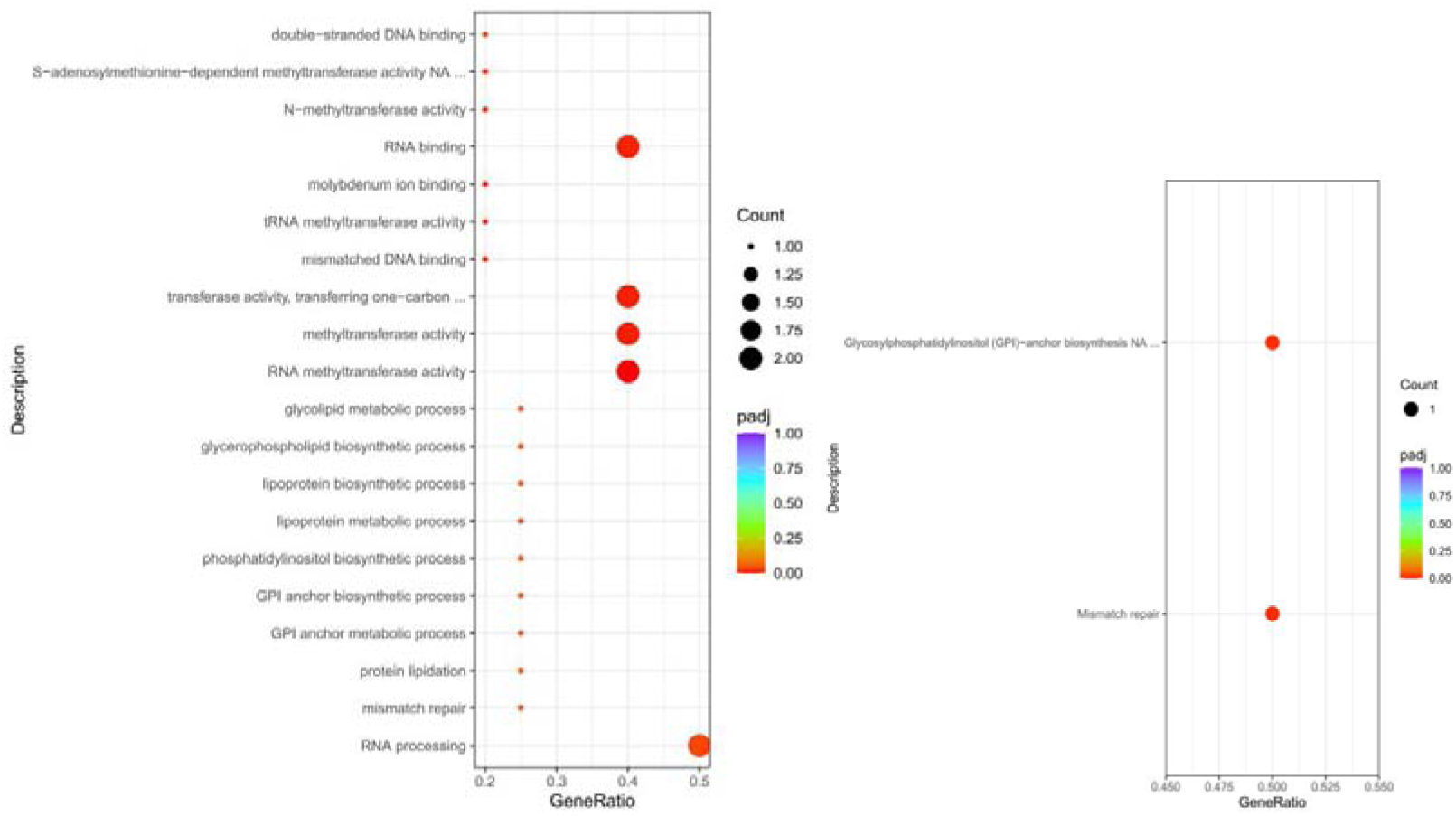
Functional enrichment of positively selected genes in the *C. chinensis*.

**Supplementary Fig.13.**
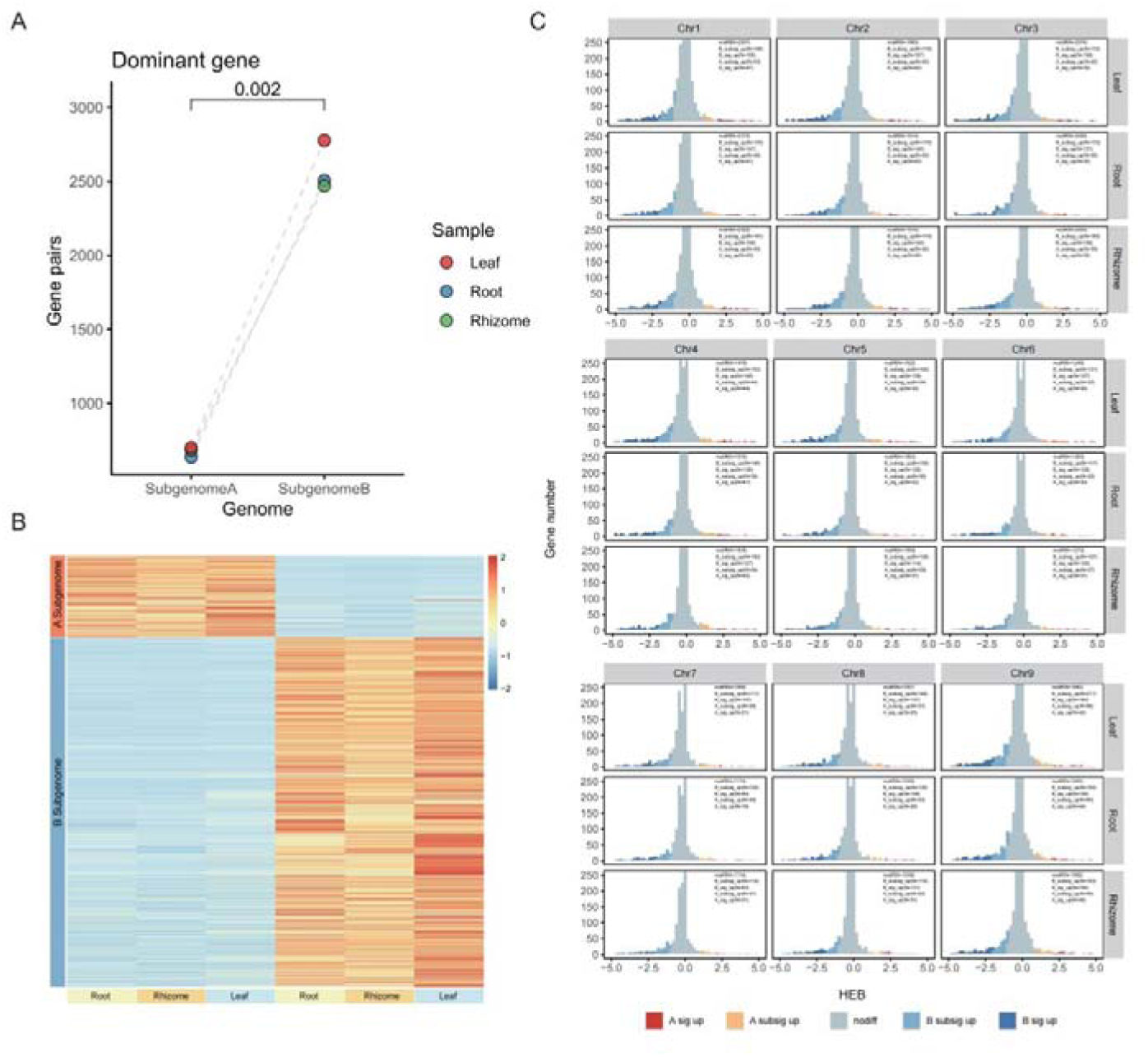
Analysis of subgenome expression dominance and homoeolog expression bias (HEB) in *C. deltoidea*.

**Supplementary Fig.14.**
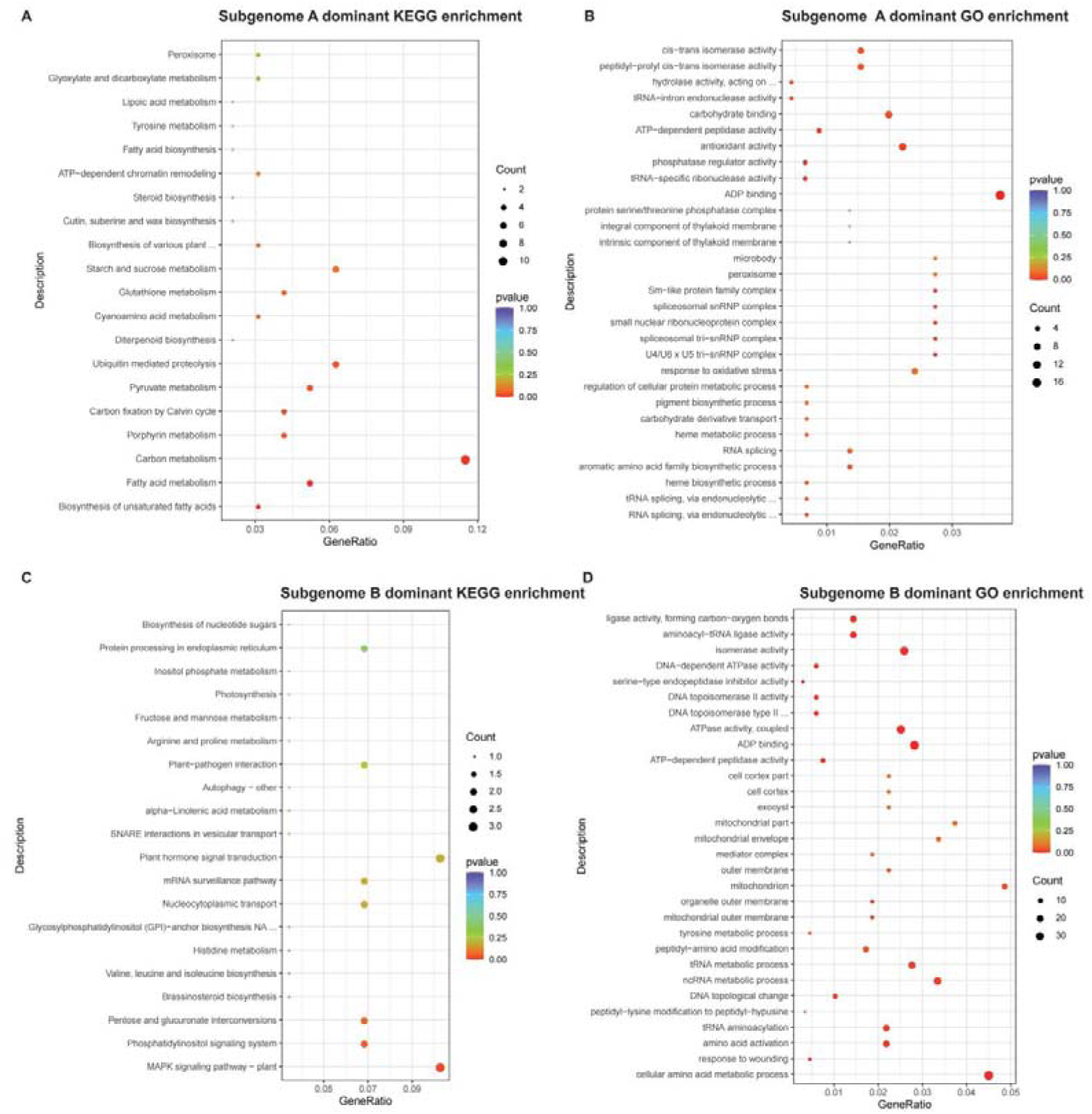
Functional enrichment of subgenome-dominant genes in *C. deltoidea*.

**Supplementary Fig.15.**
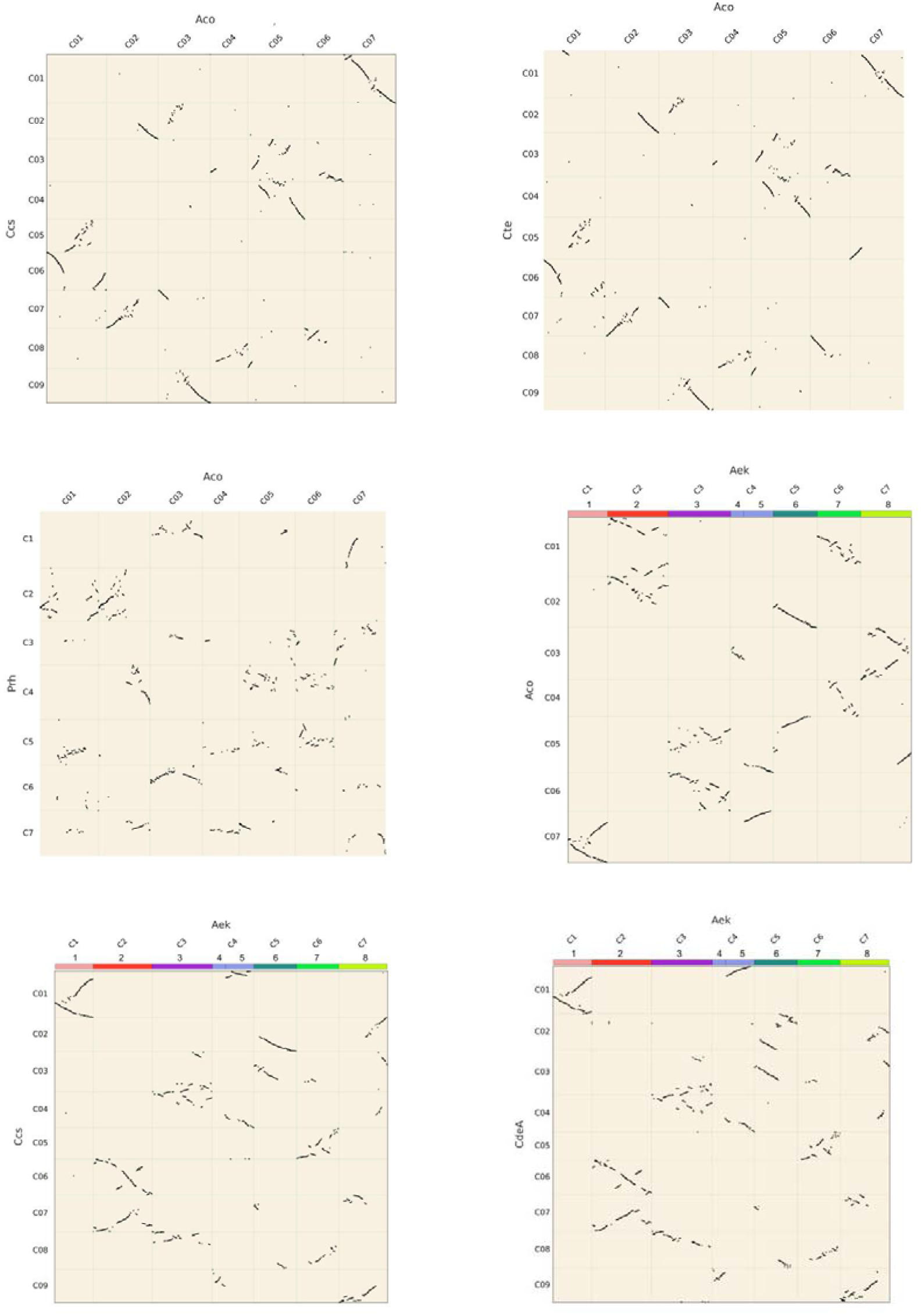
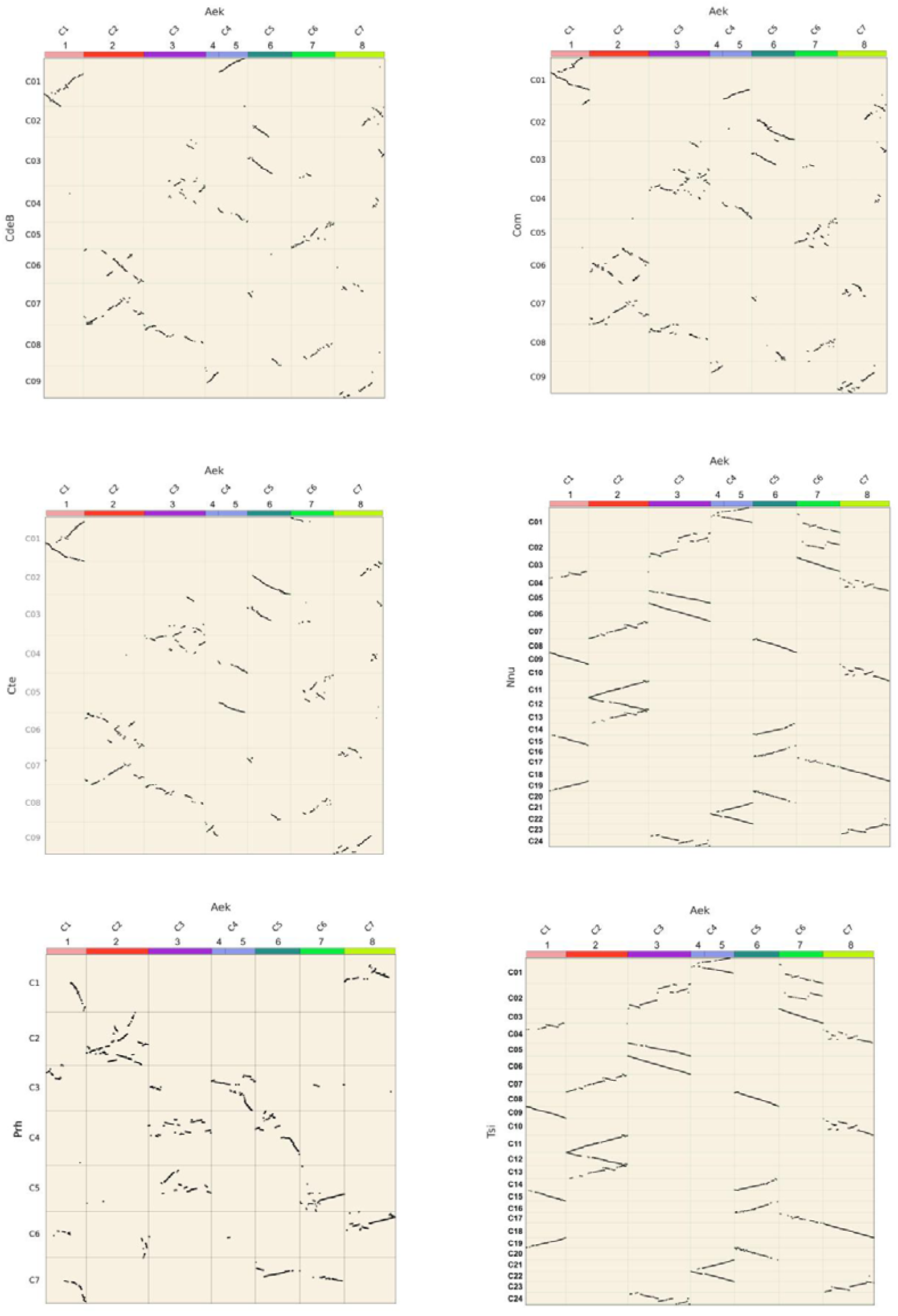
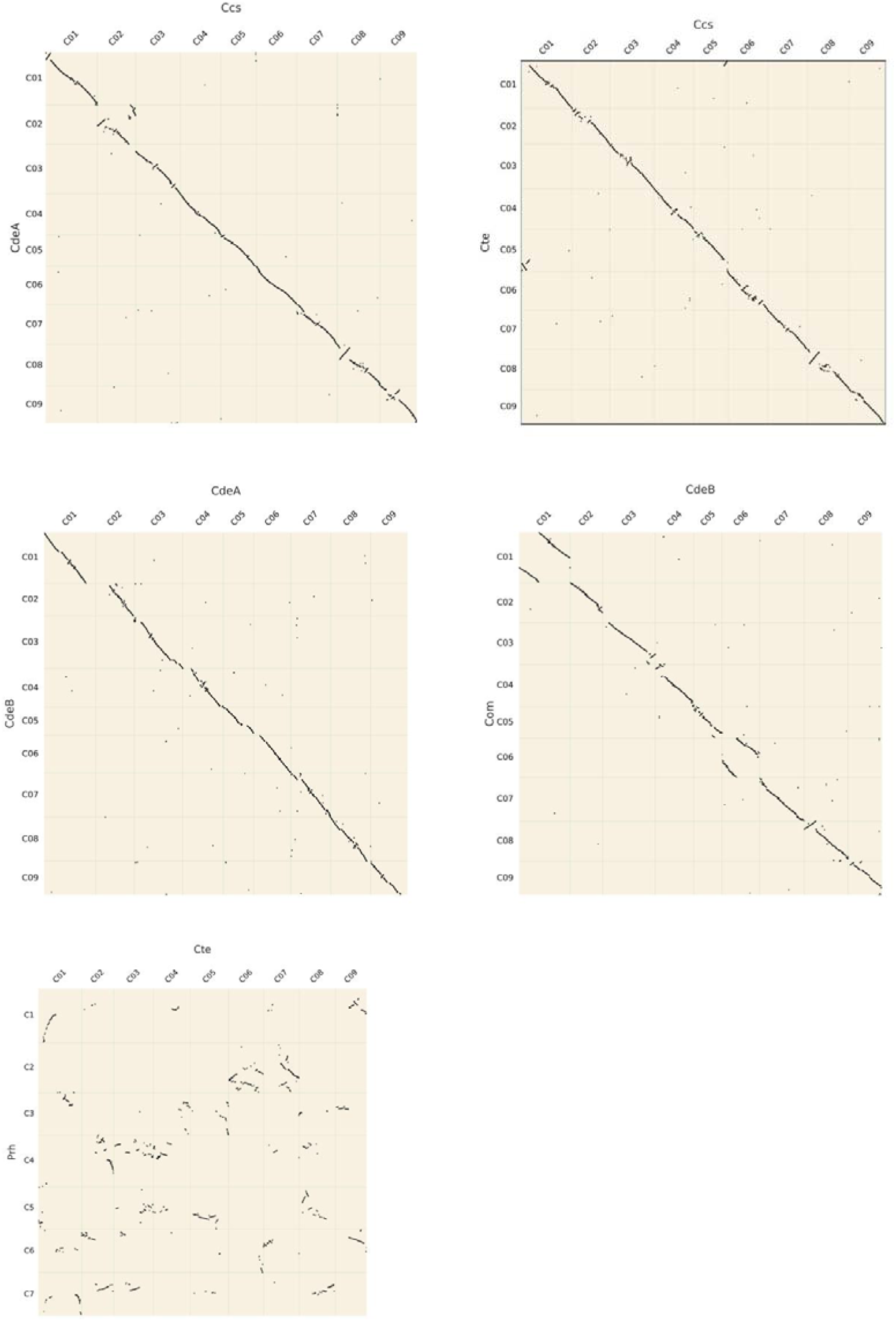
Synteny dot plots illustrating chromosomal rearrangements and conserved syntenic blocks.

**Supplementary Fig.16.**
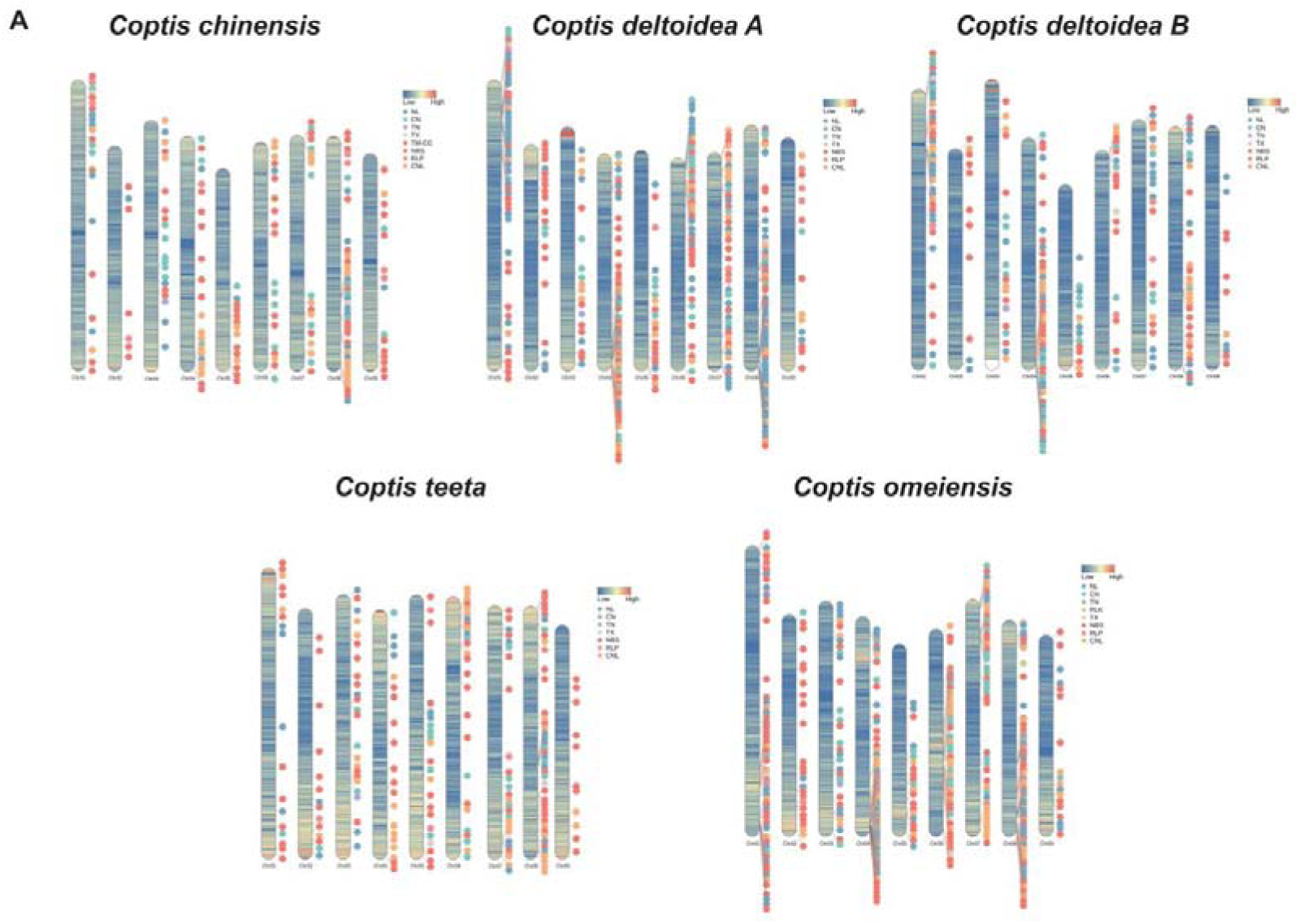
Chromosomal distribution of *NLR* genes.

**Supplementary Fig.17.**
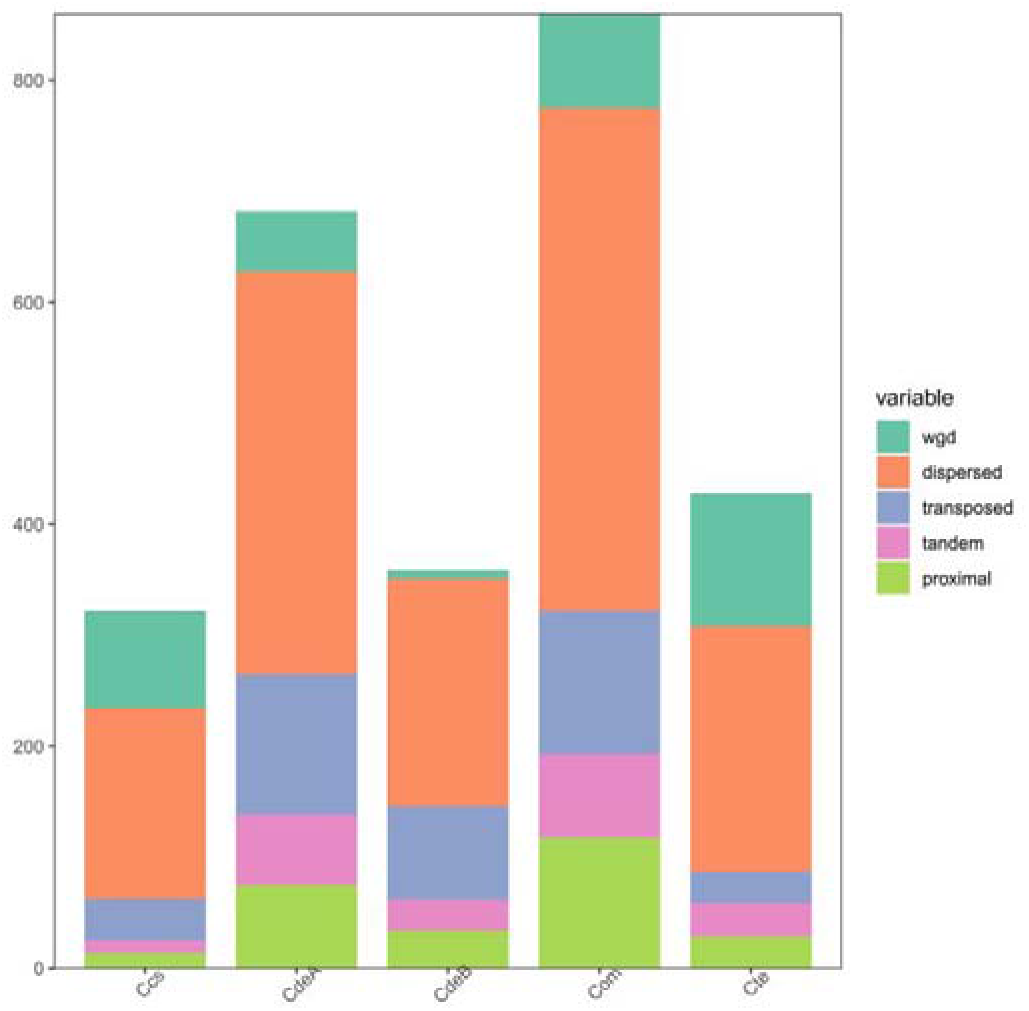
Classification of *NLR* gene duplication modes.

## References

1. Wang, J. et al. Coptidis Rhizoma: a comprehensive review of its traditional uses, botany, phytochemistry, pharmacology and toxicology. Pharm. Biol. 57, 193–225 (2019).

2. National Pharmacopoeia Commission of China. Pharmacopoeia of the People’s Republic of China: Part One (China Medical Science Press, 2020).

3. Meng, F.-C. et al. Coptidis rhizoma and its main bioactive components: recent advances in chemical investigation, quality evaluation and pharmacological activity. Chin. Med. 13, 13 (2018).

4. Li, J., Fan, G. & He, Y. Predicting the current and future distribution of three Coptis herbs in China under climate change conditions, using the MaxEnt model and chemical analysis. Sci. Total Environ. 698, 134141 (2020).

5. Pandit, M. Cytology and taxonomy of Coptis teeta Wall. (Ranunculaceae). Bot. J. Linn. Soc. 111, 371–378 (1993).

6. Cheng, Z. et al. Coptis austrogaoligongensis (Ranunculaceae), a new species from West Yunnan, China. PhytoKeys 244, 225–235 (2024).

7. Chen, H., Fan, G. & He, Y. Species evolution and quality evaluation of four Coptis herbal medicinal materials in Southwest China. 3 Biotech 7, 62 (2017).

8. Huang, J. et al. Chromosome karyotype analysis of Coptis plants in China. Acta Bot. Boreali Occident. Sin. 33, 931–938 (2013).

9. Liu, Y. et al. Analysis of the Coptis chinensis genome reveals the diversification of protoberberine-type alkaloids. Nat. Commun. 12, 3276 (2021).

10. Wang, Y. et al. A haplotype-resolved genome assembly of Coptis teeta, an endangered plant of significant medicinal value. Sci. Data 11, 1012 (2024).

11. Yang, X. et al. Three chromosome-scale Papaver genomes reveal punctuated patchwork evolution of the morphinan and noscapine biosynthesis pathway. Nat. Commun. 12, 6030 (2021).

12. An, Z. et al. Lineage specific CYP80 expansion and benzylisoquinoline alkaloid diversity in early diverging eudicots. Adv. Sci. 11, e2309990 (2024).

13. Xie, J. et al. A chromosome-scale reference genome of Aquilegia oxysepala var. kansuensis. Hortic. Res. 7, 113 (2020).

14. Ding, W.-N., Ree, R. H., Spicer, R. A. & Xing, Y.-W. Ancient orogenic and monsoon-driven assembly of the world’s richest temperate alpine flora. Science 369, 578–581 (2020).

15. Cao, G.-L. et al. Formation of three great Asian plateaus, climate change, and biodiversity. Trends Ecol. Evol. 40, 1234–1245 (2025).

16. Shimada, Y. et al. Organ-specific expression of brassinosteroid-biosynthetic genes and distribution of endogenous brassinosteroids in arabidopsis. Plant Physiol. 131, 287–297 (2003).

17. Tanaka, K. et al. Brassinosteroid homeostasis in arabidopsis is ensured by feedback expressions of multiple genes involved in its metabolism. Plant Physiol. 138, 1117–1125 (2005).

18. Jones, J. D. G., Vance, R. E. & Dangl, J. L. Intracellular innate immune surveillance devices in plants and animals. Science 354, aaf6395 (2016).

19. Shao, Z.-Q. et al. Large-scale analyses of angiosperm nucleotide-binding site-leucine-rich repeat genes reveal three anciently diverged classes with distinct evolutionary patterns. Plant Physiol. 170, 2095–2109 (2016).

20. Liu, Y. et al. An angiosperm NLR Atlas reveals that NLR gene reduction is associated with ecological specialization and signal transduction component deletion. Mol. Plant 14, 2015–2031 (2021).

21. Wang, B., Smith, S. M. & Li, J. Genetic regulation of shoot architecture. Annu. Rev. Plant Biol. 69, 437–468 (2018).

22. Shi, Z. et al. GhBZR3 suppresses cotton fiber elongation by inhibiting very long chain fatty acid biosynthesis. Plant J. 111, 785–799 (2022).

23. Ding, L. et al. Timing and mechanisms of Tibetan Plateau uplift. Nat. Rev. Earth Environ. 3, 652–667 (2022).

24. Li, Y. et al. Cenozoic India-Asia collision driven by mantle dragging the cratonic root. Nat. Commun. 15, 6674 (2024).

25. Ma, J., Song, X., Bunge, H.-P., Fichtner, A. & Tian, Y. Wholesale flat subduction of the Indian slab and northward mantle convective flow: plateau growth and driving force of the India-Asia collision. Proc. Natl Acad. Sci. U. S. A. 122, e2411776122 (2025).

26. Guo, B.-C. et al. Deciphering plant NLR genomic evolution: synteny-informed classification unveils insights into TNL gene loss. Mol. Biol. Evol. 42, msaf015 (2025).

27. Nolan, T. M., Vukašinović, N., Liu, D., Russinova, E. & Yin, Y. Brassinosteroids: multidimensional regulators of plant growth, development, and stress responses. Plant Cell 32, 295–318 (2020).

28. Han, C. et al. Brassinosteroid signaling and molecular crosstalk with nutrients in plants. J. Genet. Genomics 50, 541–553 (2023).

29. Yin, W. et al. Understanding brassinosteroid-centric phytohormone interactions for crop improvement. J. Integr. Plant Biol. 67, 563–581 (2025).

30. Ramírez-González, R. H. et al. The transcriptional landscape of polyploid wheat. Science 361, eaar6089 (2018).

31. Yoo, M.-J., Szadkowski, E. & Wendel, J. F. Homoeolog expression bias and expression level dominance in allopolyploid cotton. Heredity 110, 171–180 (2013).

32. Edger, P. P. et al. Subgenome dominance in an interspecific hybrid, synthetic allopolyploid, and a 140-year-old naturally established neo-allopolyploid monkeyflower. Plant Cell 29, 2150–2167 (2017).

33. Bird, K. A., VanBuren, R., Puzey, J. R. & Edger, P. P. Replaying the evolutionary tape to investigate subgenome dominance in allopolyploid Brassica napus. New Phytol. 230, 354–371 (2021).

34. Jarret, R. L. & Gawel, N. Abscisic acid-induced growth inhibition of sweet potato (Ipomoea batatas L.) in vitro. Plant Cell Tissue Organ Cult. 24, 13–18 (1991).

35. Belaghzal, H., Dekker, J. & Gibcus, J.H. Hi-C 2.0: an optimized Hi-C procedure for high-resolution genome-wide mapping of chromosome conformation. Methods 123, 56–65 (2017)

36. Cheng, H., Concepcion, G. T., Feng, X., Zhang, H. & Li, H. Haplotype-resolved de novo assembly using phased assembly graphs with hifiasm. Nat. Methods 18, 170–175 (2021).

37. Li, H. Aligning sequence reads, clone sequences and assembly contigs with BWA-MEM. arXiv preprint arXiv:1303.3997 (2013).

38. Durand, N. C. et al. Juicer provides a one-click system for analyzing loop-resolution Hi-C experiments. Cell Syst. 3, 95–98 (2016).

39. Zheng, J. et al. From molecular to macroscopic via the rational design of a self-assembled 3D DNA crystal. Nature 461, 74–77 (2009).

40. Durand, N. C. et al. Juicebox provides a visualization system for Hi-C contact maps with unlimited zoom. Cell Syst. 3, 99–101 (2016).

41. Manni, M., Berkeley, M. R., Seppey, M., Simão, F. A. & Zdobnov, E. M. BUSCO update: novel and streamlined workflows along with broader and deeper phylogenetic coverage for scoring of eukaryotic, prokaryotic, and viral genomes. Mol. Biol. Evol. 38, 4647–4654 (2021).

42. Ou, S., Chen, J. & Jiang, N. Assessing genome assembly quality using the LTR Assembly Index (LAI). Nucleic Acids Res. 46, e126 (2018).

43. Moereels, H., De Bie, L. & Tollenaere, J. P. CGEMA and VGAP: a colour graphics editor for multiple alignment using a variable GAP penalty. Application to the muscarinic acetylcholine receptor. J. Comput. Aided Mol. Des. 4, 131-145 (1990).

44. Flynn, J. M. et al. RepeatModeler2 for automated genomic discovery of transposable element families. Proc. Natl Acad. Sci. U. S. A. 117, 9451–9457 (2020).

45. Xu, Z. & Wang, H. LTR_FINDER: an efficient tool for the prediction of full-length LTR retrotransposons. Nucleic Acids Res. 35, W265–W268 (2007).

46. Chen, N. Using RepeatMasker to identify repetitive elements in genomic sequences. Curr. Protoc. Bioinform. 5, 4.10.11–4.10.14 (2004).

47. Benson, G. Tandem repeats finder: a program to analyze DNA sequences. Nucleic Acids Res. 27, 573–580 (1999).

48. Stanke, M. et al. AUGUSTUS: ab initio prediction of alternative transcripts. Nucleic Acids Res. 34, W435–W439 (2006).

49. Johnson, A. D. et al. SNAP: a web-based tool for identification and annotation of proxy SNPs using HapMap. Bioinformatics 24, 2938–2939 (2008).

50. Keilwagen, J., Hartung, F. & Grau, J. GeMoMa: homology-based gene prediction utilizing intron position conservation and RNA-seq data in Gene Prediction: Methods and Protocols (ed. Kollmar, M.) 161-177 (Springer, 2019).

51. Grabherr, M. G. et al. Trinity: reconstructing a full-length transcriptome without a genome from RNA-Seq data. Nat. Biotechnol. 29, 644–652 (2011).

52. Haas, B. J. et al. Automated eukaryotic gene structure annotation using EVidenceModeler and the Program to Assemble Spliced Alignments. Genome Biol. 9, R7 (2008).

53. Wingett, S. W. & Andrews, S. FastQ screen: a tool for multi-genome mapping and quality control. F1000Research 7, 1338 (2018).

54. Bolger, A. M., Lohse, M. & Usadel, B. Trimmomatic: a flexible trimmer for Illumina sequence data. Bioinformatics 30, 2114–2120 (2014).

55. Dobin, A. et al. STAR: ultrafast universal RNA-seq aligner. Bioinformatics 29, 15–21 (2012).

56. Li, H. et al. The sequence alignment/map format and SAMtools. Bioinformatics 25, 2078–2079 (2009).

57. Pertea, M. et al. StringTie enables improved reconstruction of a transcriptome from RNA-seq reads. Nat. Biotechnol. 33, 290–295 (2015).

58. Pertea, G. & Pertea, M. GFF utilities: GffRead and GffCompare. F1000Research 9, 304 (2020).

59. Emms, D. M. & Kelly, S. OrthoFinder: phylogenetic orthology inference for comparative genomics. Genome Biol. 20, 238 (2019).

60. Buchfink, B., Xie, C. & Huson, D. H. Fast and sensitive protein alignment using DIAMOND. Nat. Methods 12, 59–60 (2015).

61. Katoh, K. MAFFT: a novel method for rapid multiple sequence alignment based on fast Fourier transform. Nucleic Acids Res. 30, 3059–3066 (2002).

62. Yang, Z. & Rannala, B. Bayesian estimation of species divergence times under a molecular clock using multiple fossil calibrations with soft bounds. Mol. Biol. Evol. 23, 212–226 (2006).

63. Kumar, S., Stecher, G., Suleski, M. & Hedges, S. B. TimeTree: a resource for timelines, timetrees, and divergence times. Mol. Biol. Evol. 34, 1812–1819 (2017).

64. Letunic, I. & Bork, P. Interactive tree of life (iTOL) v5: an online tool for phylogenetic tree display and annotation. Nucleic Acids Res. 49, W293–W296 (2021).

65. Mendes, F. K., Vanderpool, D., Fulton, B. & Hahn, M. W. CAFE 5 models variation in evolutionary rates among gene families. Bioinformatics 36, 5516–5518 (2020).

66. Yang, Z. PAML 4: phylogenetic analysis by maximum likelihood. Mol. Biol. Evol. 24, 1586–1591 (2007).

67. Wang, Y. et al. MCScanX: a toolkit for detection and evolutionary analysis of gene synteny and collinearity. Nucleic Acids Res. 40, e49 (2012).

68. Pham, S. K. & Pevzner, P. A. DRIMM-Synteny: decomposing genomes into evolutionary conserved segments. Bioinformatics 26, 2509–2516 (2010).

69. Gao, S. et al. IAGS: inferring ancestor genome structure under a wide range of evolutionary scenarios. Mol. Biol. Evol. 39, msac041 (2022).

70. Jia, K. H. et al. SubPhaser: a robust allopolyploid subgenome phasing method based on subgenome specific k mers. New Phytol. 235, 801–809 (2022).

71. Li, P. et al. RGAugury: a pipeline for genome-wide prediction of resistance gene analogs (RGAs) in plants. BMC Genom. 17, 852 (2016).

72. Nguyen, L.-T., Schmidt, H. A., von Haeseler, A. & Minh, B. Q. IQ-TREE: a fast and effective stochastic algorithm for estimating maximum-likelihood phylogenies. Mol. Biol. Evol. 32, 268–274 (2015).

73. Qiao, X. et al. Gene duplication and evolution in recurring polyploidization–diploidization cycles in plants. Genome Biol. 20, 38 (2019).

74. Wu, L., et al. A biosynthetic network for protoberberine production in Coptis chinensis. Horticulture Res. 11, uhad259 (2024).

75. Chen, D. et al. The chromosome-level reference genome of Coptis chinensis provides insights into genomic evolution and berberine biosynthesis. Horticulture Res. 8, 1 (2021).

76. Langfelder, P. & Horvath, S. WGCNA: an R package for weighted correlation network analysis. BMC Bioinformatics 9, 559 (2008).

77. Shannon, P. et al. Cytoscape: a software environment for integrated models of biomolecular interaction networks. Genome Res. 13, 2498–2504 (2003).

78. Namuag, P. MySQL Performance Benchmarking: MySQL 5.7 vs MySQL 8.0 (2019).

79. Sun, J. et al. OrthoVenn3: an integrated platform for exploring and visualizing orthologous data across genomes. Nucleic Acids Res. 51, W397–W403 (2023).

80. Zheng, Q. & Wang, X.-J. GOEAST: a web-based software toolkit for Gene Ontology enrichment analysis. Nucleic Acids Res. 36, W358–W363 (2008).

81. Li, D. et al. ECharts: a declarative framework for rapid construction of web-based visualization. Vis. Inform. 2, 136–146 (2018).

82. Szmulewicz, D. J. et al. CANVAS an update: clinical presentation, investigation and management. J. Vestib. Res. 24, 465–474 (2014).

